## Supplemental Materials for "Divergent Folding-Mediated Epistasis Among Unstable Membrane Protein Variants"

**This Document Includes:**

-Figure S1
-Figure S2
-Figure S3
-Table S1
-Table S2
-Table S3

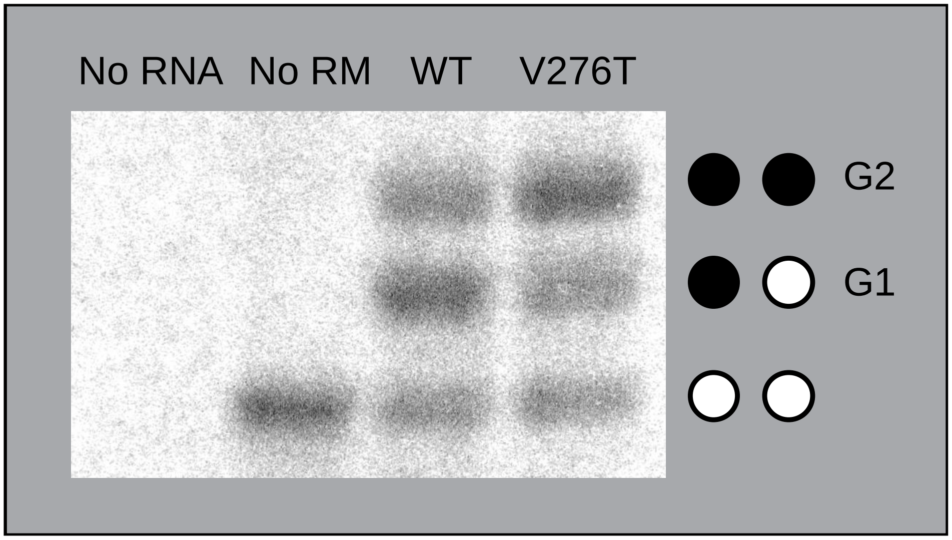


**Figure S1. Membrane integration of mGnRHR TM domain 6 and V276T.** Chimeric Lep proteins containing either the wild type mGnRHR TM domain 6 or the V276T mutant, with the guest TM domain flanked by two glycosylation sites, were translated *in vitro* and analyzed by SDS-PAGE. Misintegration of the guest TM domain results in doubly glycosylated protein, while successful membrane integration results in singly glycosylated protein. Negative control translation reactions without RNA (first lane) and without rough microsomes (second lane) are included. The positions of the untargeted protein that is not associated with the membrane (no glycosylation), singly glycosylated form of the protein in which TMD6 is integrated into the membrane (G1), and doubly glycosylated form of the protein in which TMD6 is not integrated into the membrane (G2) are indicated for reference.

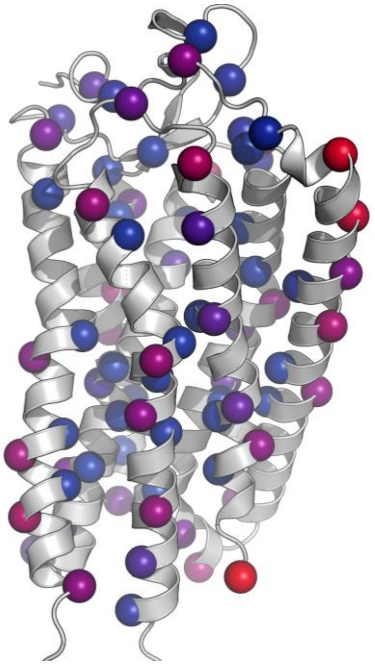

**Figure S2. Structural context of mGnRHR library mutants.** Residues selected for mutation are displayed as spheres on the homology model of *M. musculus* GnRHR. Spheres are colored according to solvent accessible surface area of the residue, with red representing exposed residues and blue representing buried residues.

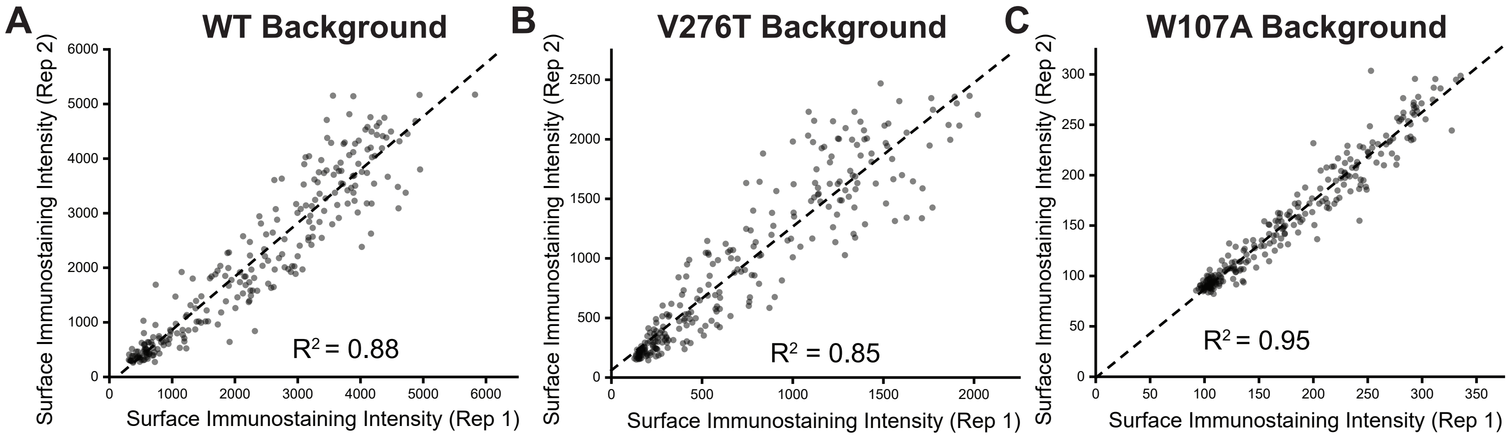


**Figure S3. Correlation of DMS Variant Intensity Values Across Two Biological Replicates.** The inten-sity values for the surface immunostaining of the 251 mutants scored the A) WT, B) V276T, and C) W107A backgrounds determined from two independent biological replicates are plotted against one another. A linear fit (dashes) is shown for reference along with the Pearson's $R^2$ value, for reference. These data demonstrate the precision of these measurements despite variations in the signal-to noise that arises from variations in surface immunostaining intensity.

**Table S1. Predicted and measured apparent transfer free energies of mGnRHR TM6.**

| **Insert TMD** | **$K_{app}$ ‡<br>(Int. $_{G1}$ / Int. $_{G2}$)** | **$\Delta G_{app}$ ‡<br>(kcal/ mol)** | **$\Delta G_{app, pred}$ †<br>(kcal/ mol)** |
|---|---|---|---|
| WT TMD6 | 1.2 ± 0.2 | -0.1 ± 0.1 | 0.26 |
| V276T TMD6 | 0.7 ± 0.1 | 0.2 ± 0.1 | 0.805 |

‡Values represent the average of three experimental replicates, and errors represent the standard deviation.
†Values predicted from the TM6 sequence using the ΔG Predictor.

**Table S2. Solvent accessible surface area of mGnRHR residues.**

| Position | Residue | Percent Relative Accessibility (REL) |
|---|---|---|
| 13 | His | 91.1 |
| 14 | Cys | 63.9 |
| 15 | Ser | 21.9 |
| 16 | Ala | 32.1 |
| 17 | Ile | 67.3 |
| 18 | Asn | 76.5 |
| 19 | Asn | 1.3 |
| 20 | Ser | 15.6 |
| 21 | Ile | 3.9 |
| 22 | Pro | 27.2 |
| 23 | Leu | 3.2 |
| 24 | Ile | 50.7 |
| 25 | Gln | 33.3 |
| 26 | Gly | 9.0 |
| 27 | Lys | 69.8 |
| 28 | Leu | 7.6 |
| 29 | Pro | 44.6 |
| 30 | Thr | 45.0 |
| 31 | Leu | 13.5 |
| 32 | Thr | 84.7 |
| 33 | Val | 69.0 |
| 34 | Ser | 13.8 |
| 35 | Gly | 4.5 |
| 36 | Lys | 80.6 |
| 37 | Ile | 51.7 |
| 38 | Arg | 7.0 |
| 39 | Val | 29.3 |
| 40 | Thr | 49.0 |
| 41 | Val | 31.7 |
| 42 | Thr | 12.3 |
| 43 | Phe | 59.6 |
| 44 | Phe | 62.2 |
| 45 | Leu | 3.5 |
| 46 | Phe | 27.5 |
| 47 | Leu | 64.6 |
| 48 | Leu | 31.0 |
| 49 | Ser | 0.0 |
| 50 | Thr | 31.0 |

| | | |
|---|---|---|
| 51 | Ala | 51.6 |
| 52 | Phe | 29.7 |
| 53 | Asn | 0.0 |
| 54 | Ala | 30.6 |
| 55 | Ser | 40.6 |
| 56 | Phe | 7.0 |
| 57 | Leu | 11.6 |
| 58 | Leu | 57.0 |
| 59 | Lys | 54.6 |
| 60 | Leu | 0.7 |
| 61 | Gln | 44.7 |
| 62 | Lys | 72.6 |
| 63 | Trp | 26.1 |
| 64 | Thr | 49.3 |
| 65 | Gln | 81.5 |
| 66 | Lys | 53.8 |
| 67 | Arg | 91.8 |
| 68 | Lys | 59.2 |
| 69 | Lys | 34.7 |
| 70 | Gly | 24.7 |
| 71 | Lys | 71.5 |
| 72 | Lys | 42.9 |
| 73 | Leu | 3.7 |
| 74 | Ser | 10.8 |
| 75 | Arg | 18.0 |
| 76 | Met | 4.0 |
| 77 | Lys | 7.5 |
| 78 | Val | 9.2 |
| 79 | Leu | 0.1 |
| 80 | Leu | 2.2 |
| 81 | Lys | 21.5 |
| 82 | His | 14.5 |
| 83 | Leu | 1.5 |
| 84 | Thr | 0.0 |
| 85 | Leu | 33.8 |
| 86 | Ala | 0.2 |
| 87 | Asn | 6.0 |
| 88 | Leu | 17.4 |
| 89 | Leu | 37.1 |
| 90 | Glu | 4.4 |
| 91 | Thr | 0.1 |

| | | |
|---|---|---|
| 92 | Leu | 52.1 |
| 93 | Ile | 44.3 |
| 94 | Val | 3.0 |
| 95 | Met | 3.0 |
| 96 | Pro | 49.0 |
| 97 | Leu | 16.9 |
| 98 | Asp | 3.0 |
| 99 | Gly | 16.4 |
| 100 | Met | 27.2 |
| 101 | Trp | 14.8 |
| 102 | Asn | 5.0 |
| 103 | Ile | 21.9 |
| 104 | Thr | 43.7 |
| 105 | Val | 5.6 |
| 106 | Gln | 12.7 |
| 107 | Trp | 0.5 |
| 108 | Tyr | 57.0 |
| 109 | Ala | 17.3 |
| 110 | Gly | 49.6 |
| 111 | Glu | 36.5 |
| 112 | Phe | 69.9 |
| 113 | Leu | 34.1 |
| 114 | Cys | 0.2 |
| 115 | Lys | 21.2 |
| 116 | Val | 11.2 |
| 117 | Leu | 9.3 |
| 118 | Ser | 14.0 |
| 119 | Tyr | 24.0 |
| 120 | Leu | 30.6 |
| 121 | Lys | 3.3 |
| 122 | Leu | 5.6 |
| 123 | Phe | 23.2 |
| 124 | Ser | 6.1 |
| 125 | Met | 6.5 |
| 126 | Tyr | 0.0 |
| 127 | Ala | 0.0 |
| 128 | Pro | 7.3 |
| 129 | Ala | 1.3 |
| 130 | Phe | 21.6 |
| 131 | Met | 0.3 |
| 132 | Met | 5.8 |

| | | |
|---|---|---|
| 133 | Val | 11.5 |
| 134 | Val | 6.7 |
| 135 | Ile | 2.4 |
| 136 | Ser | 0.0 |
| 137 | Leu | 43.5 |
| 138 | Asp | 2.1 |
| 139 | Arg | 9.4 |
| 140 | Ser | 22.0 |
| 141 | Leu | 21.4 |
| 142 | Ala | 15.6 |
| 143 | Ile | 14.6 |
| 144 | Thr | 31.0 |
| 145 | Gln | 59.5 |
| 146 | Pro | 50.1 |
| 147 | Leu | 95.7 |
| 148 | Ala | 35.3 |
| 149 | Val | 67.0 |
| 150 | Gln | 65.6 |
| 151 | Ser | 4.0 |
| 152 | Asn | 41.5 |
| 153 | Ser | 40.8 |
| 154 | Lys | 85.6 |
| 155 | Leu | 40.3 |
| 156 | Glu | 2.4 |
| 157 | Glm | 40.2 |
| 158 | Ser | 41.0 |
| 159 | Met | 33.0 |
| 160 | Ile | 0.8 |
| 161 | Ser | 45.5 |
| 162 | Leu | 46.4 |
| 163 | Ala | 1.5 |
| 164 | Trp | 17.8 |
| 165 | Ile | 52.0 |
| 166 | Leu | 37.9 |
| 167 | Ser | 0.0 |
| 168 | Ile | 60.8 |
| 169 | Val | 58.1 |
| 170 | Phe | 24.9 |
| 171 | Ala | 4.4 |
| 172 | Gly | 45.4 |
| 173 | Pro | 23.8 |

| | | |
|---|---|---|
| 174 | Gln | 3.7 |
| 175 | Leu | 40.7 |
| 176 | Tyr | 29.5 |
| 177 | Ile | 0.0 |
| 178 | Phe | 13.1 |
| 179 | Arg | 36.4 |
| 180 | Met | 21.9 |
| 181 | Ile | 7.9 |
| 182 | Tyr | 64.6 |
| 183 | Leu | 40.2 |
| 184 | Ala | 0.3 |
| 185 | Asp | 20.1 |
| 186 | Gly | 97.0 |
| 187 | Ser | 26.6 |
| 188 | Gly | 2.2 |
| 189 | Pro | 80.5 |
| 190 | Thr | 70.8 |
| 191 | Val | 63.0 |
| 192 | Phe | 11.8 |
| 193 | Ser | 2.2 |
| 194 | Gln | 5.3 |
| 195 | Cys | 9.0 |
| 196 | Val | 6.3 |
| 197 | Thr | 0.5 |
| 198 | His | 4.9 |
| 199 | Cys | 0.0 |
| 200 | Ser | 21.3 |
| 201 | Phe | 54.6 |
| 202 | Pro | 38.7 |
| 203 | Gln | 40.4 |
| 204 | Trp | 43.1 |
| 205 | Trp | 43.3 |
| 206 | His | 19.3 |
| 207 | Gln | 0.0 |
| 208 | Ala | 3.9 |
| 209 | Phe | 31.8 |
| 210 | Tyr | 26.2 |
| 211 | Asn | 0.7 |
| 212 | Phe | 57.2 |
| 213 | Phe | 46.1 |
| 214 | Thr | 5.6 |

| | | |
|---|---|---|
| 215 | Phe | 19.7 |
| 216 | Gly | 19.0 |
| 217 | Cys | 24.8 |
| 218 | Leu | 4.1 |
| 219 | Phe | 6.3 |
| 220 | Ile | 34.9 |
| 221 | Ile | 50.2 |
| 222 | Pro | 7.7 |
| 223 | Leu | 20.8 |
| 224 | Leu | 52.4 |
| 225 | Ile | 36.8 |
| 226 | Met | 2.7 |
| 227 | Leu | 55.4 |
| 228 | Ile | 58.2 |
| 229 | Cys | 14.9 |
| 230 | Asn | 7.5 |
| 231 | Ala | 35.5 |
| 232 | Lys | 63.7 |
| 233 | Ile | 0.1 |
| 234 | Ile | 34.0 |
| 235 | Phe | 70.0 |
| 236 | Ala | 25.7 |
| 237 | Leu | 20.4 |
| 238 | Thr | 51.5 |
| 239 | Arg | 73.4 |
| 240 | Val | 42.9 |
| 241 | Leu | 74.1 |
| 242 | His | 96.1 |
| 243 | Gln | 64.9 |
| 244 | Asp | 23.5 |
| 245 | Pro | 44.3 |
| 246 | Arg | 67.7 |
| 247 | Lys | 51.7 |
| 248 | Leu | 41.1 |
| 249 | Gln | 46.3 |
| 250 | Leu | 51.1 |
| 251 | Asn | 26.2 |
| 252 | Gln | 76.8 |
| 253 | Ser | 71.5 |
| 254 | Lys | 61.6 |
| 255 | Asn | 93.2 |

| | | |
|---|---|---|
| 256 | Asn | 43.8 |
| 257 | Ile | 55.2 |
| 258 | Pro | 12.6 |
| 259 | Arg | 72.2 |
| 260 | Ala | 39.4 |
| 261 | Arg | 29.7 |
| 262 | Leu | 31.1 |
| 263 | Arg | 8.3 |
| 264 | Thr | 13.3 |
| 265 | Leu | 4.2 |
| 266 | Lys | 55.6 |
| 267 | Met | 2.4 |
| 268 | Thr | 5.2 |
| 269 | Val | 39.8 |
| 270 | Ala | 39.5 |
| 271 | Phe | 3.3 |
| 272 | Ala | 13.6 |
| 273 | Thr | 51.0 |
| 274 | Ser | 9.2 |
| 275 | Phe | 6.0 |
| 276 | Val | 28.4 |
| 277 | Val | 57.2 |
| 278 | Cys | 1.4 |
| 279 | Trp | 2.4 |
| 280 | Thr | 17.6 |
| 281 | Pro | 20.2 |
| 282 | Tyr | 14.6 |
| 283 | Tyr | 6.5 |
| 284 | Val | 29.8 |
| 285 | Leu | 3.1 |
| 286 | Gly | 7.2 |
| 287 | Ile | 26.7 |
| 288 | Trp | 56.9 |
| 289 | Tyr | 18.0 |
| 290 | Trp | 23.1 |
| 291 | Phe | 35.9 |
| 292 | Asp | 51.7 |
| 293 | Pro | 14.0 |
| 294 | Glu | 81.8 |
| 295 | Met | 41.9 |
| 296 | Leu | 11.8 |

| | | |
|---|---|---|
| 297 | Asn | 53.7 |
| 298 | Arg | 48.4 |
| 299 | Val | 65.5 |
| 300 | Ser | 16.8 |
| 301 | Glu | 3.0 |
| 302 | Pro | 17.3 |
| 303 | Val | 34.4 |
| 304 | Asn | 4.1 |
| 305 | His | 2.9 |
| 306 | Phe | 48.8 |
| 307 | Phe | 24.6 |
| 308 | Phe | 6.8 |
| 309 | Leu | 11.4 |
| 310 | Phe | 36.9 |
| 311 | Ala | 0.1 |
| 312 | Phe | 1.9 |
| 313 | Leu | 28.8 |
| 314 | Asn | 1.3 |
| 315 | Pro | 1.2 |
| 316 | Cys | 2.2 |
| 317 | Phe | 36.7 |
| 318 | Asp | 1.2 |
| 319 | Pro | 4.4 |
| 320 | Leu | 52.2 |
| 321 | Ile | 18.7 |
| 322 | Tyr | 0.8 |
| 323 | Gly | 31.6 |
| 324 | Tyr | 66.1 |
| 325 | Phe | 26.3 |
| 326 | Ser | 62.2 |
| 327 | Leu | 89.8 |

**Table S3. mGnRHR library mutants with associated oligo sequences.**

| Mutation | Oligo Sequence |
|---|---|
| N4A | GGTACCATGGCTAACGCCGCATCTCTTGAGCAG |
| N4R | GGTACCATGGCTAACAGAGCATCTCTTGAGCAG |
| N4D | GGTACCATGGCTAACGACGCATCTCTTGAGCAG |
| N4C | GGTACCATGGCTAACTGCGCATCTCTTGAGCAG |
| N4Q | GGTACCATGGCTAACCAGGCATCTCTTGAGCAG |
| N4E | GGTACCATGGCTAACGAGGCATCTCTTGAGCAG |
| N4G | GGTACCATGGCTAACGGCGCATCTCTTGAGCAG |
| N4H | GGTACCATGGCTAACCACGCATCTCTTGAGCAG |
| N4I | GGTACCATGGCTAACATCGCATCTCTTGAGCAG |
| N4L | GGTACCATGGCTAACCTGGCATCTCTTGAGCAG |
| N4K | GGTACCATGGCTAACAAGGCATCTCTTGAGCAG |
| N4M | GGTACCATGGCTAACATGGCATCTCTTGAGCAG |
| N4F | GGTACCATGGCTAACTTCGCATCTCTTGAGCAG |
| N4P | GGTACCATGGCTAACCCCGCATCTCTTGAGCAG |
| N4S | GGTACCATGGCTAACAGCGCATCTCTTGAGCAG |
| N4T | GGTACCATGGCTAACACCGCATCTCTTGAGCAG |
| N4W | GGTACCATGGCTAACTGGGCATCTCTTGAGCAG |
| N4Y | GGTACCATGGCTAACTACGCATCTCTTGAGCAG |
| N4V | GGTACCATGGCTAACGTGGCATCTCTTGAGCAG |
| E8A | AACAATGCATCTCTTGCCCAGGACCCAAATCAC |
| E8R | AACAATGCATCTCTTAGACAGGACCCAAATCAC |
| E8N | AACAATGCATCTCTTAACCAGGACCCAAATCAC |
| E8D | AACAATGCATCTCTTGACCAGGACCCAAATCAC |
| E8C | AACAATGCATCTCTTTGCCAGGACCCAAATCAC |
| E8Q | AACAATGCATCTCTTCAGCAGGACCCAAATCAC |
| E8G | AACAATGCATCTCTTGGCCAGGACCCAAATCAC |
| E8H | AACAATGCATCTCTTCACCAGGACCCAAATCAC |
| E8I | AACAATGCATCTCTTATCCAGGACCCAAATCAC |
| E8L | AACAATGCATCTCTTCTGCAGGACCCAAATCAC |
| E8K | AACAATGCATCTCTTAAGCAGGACCCAAATCAC |
| E8M | AACAATGCATCTCTTATGCAGGACCCAAATCAC |
| E8F | AACAATGCATCTCTTTTCCAGGACCCAAATCAC |
| E8P | AACAATGCATCTCTTCCCCAGGACCCAAATCAC |
| E8S | AACAATGCATCTCTTAGCCAGGACCCAAATCAC |
| E8T | AACAATGCATCTCTTACCCAGGACCCAAATCAC |
| E8W | AACAATGCATCTCTTTGGCAGGACCCAAATCAC |
| E8Y | AACAATGCATCTCTTTACCAGGACCCAAATCAC |
| E8V | AACAATGCATCTCTTGTGCAGGACCCAAATCAC |
| N12A | CTTGAGCAGGACCCAGCCCACTGCTCGGCCATC |

| | |
|---|---|
| N12R | CTTGAGCAGGACCCAAGACACTGCTCGGCCATC |
| N12D | CTTGAGCAGGACCCAGACCACTGCTCGGCCATC |
| N12C | CTTGAGCAGGACCCATGCCACTGCTCGGCCATC |
| N12Q | CTTGAGCAGGACCCACAGCACTGCTCGGCCATC |
| N12E | CTTGAGCAGGACCCAGAGCACTGCTCGGCCATC |
| N12G | CTTGAGCAGGACCCAGGCCACTGCTCGGCCATC |
| N12H | CTTGAGCAGGACCCACACCACTGCTCGGCCATC |
| N12I | CTTGAGCAGGACCCAATCCACTGCTCGGCCATC |
| N12L | CTTGAGCAGGACCCACTGCACTGCTCGGCCATC |
| N12K | CTTGAGCAGGACCCAAAGCACTGCTCGGCCATC |
| N12M | CTTGAGCAGGACCCAATGCACTGCTCGGCCATC |
| N12F | CTTGAGCAGGACCCATTCCACTGCTCGGCCATC |
| N12P | CTTGAGCAGGACCCACCCCACTGCTCGGCCATC |
| N12S | CTTGAGCAGGACCCAAGCCACTGCTCGGCCATC |
| N12T | CTTGAGCAGGACCCAACCCACTGCTCGGCCATC |
| N12W | CTTGAGCAGGACCCATGGCACTGCTCGGCCATC |
| N12Y | CTTGAGCAGGACCCATACCACTGCTCGGCCATC |
| N12V | CTTGAGCAGGACCCAGTGCACTGCTCGGCCATC |
| A16R | CCAAATCACTGCTCGAGAATCAACAACAGCATC |
| A16N | CCAAATCACTGCTCGAACATCAACAACAGCATC |
| A16D | CCAAATCACTGCTCGGACATCAACAACAGCATC |
| A16C | CCAAATCACTGCTCGTGCATCAACAACAGCATC |
| A16Q | CCAAATCACTGCTCGCAGATCAACAACAGCATC |
| A16E | CCAAATCACTGCTCGGAGATCAACAACAGCATC |
| A16G | CCAAATCACTGCTCGGGCATCAACAACAGCATC |
| A16H | CCAAATCACTGCTCGCACATCAACAACAGCATC |
| A16I | CCAAATCACTGCTCGATCATCAACAACAGCATC |
| A16L | CCAAATCACTGCTCGCTGATCAACAACAGCATC |
| A16K | CCAAATCACTGCTCGAAGATCAACAACAGCATC |
| A16M | CCAAATCACTGCTCGATGATCAACAACAGCATC |
| A16F | CCAAATCACTGCTCGTTCATCAACAACAGCATC |
| A16P | CCAAATCACTGCTCGCCCATCAACAACAGCATC |
| A16S | CCAAATCACTGCTCGAGCATCAACAACAGCATC |
| A16T | CCAAATCACTGCTCGACCATCAACAACAGCATC |
| A16W | CCAAATCACTGCTCGTGGATCAACAACAGCATC |
| A16Y | CCAAATCACTGCTCGTACATCAACAACAGCATC |
| A16V | CCAAATCACTGCTCGGTGATCAACAACAGCATC |
| S20A | TCGGCCATCAACAACGCCATCCCCTTGATACAG |
| S20R | TCGGCCATCAACAACAGAATCCCCTTGATACAG |
| S20N | TCGGCCATCAACAACAACATCCCCTTGATACAG |
| S20D | TCGGCCATCAACAACGACATCCCCTTGATACAG |

| | |
|---|---|
| S20C | TCGGCCATCAACAACTGCATCCCCTTGATACAG |
| S20Q | TCGGCCATCAACAACCAGATCCCCTTGATACAG |
| S20E | TCGGCCATCAACAACGAGATCCCCTTGATACAG |
| S20G | TCGGCCATCAACAACGGCATCCCCTTGATACAG |
| S20H | TCGGCCATCAACAACCACATCCCCTTGATACAG |
| S20I | TCGGCCATCAACAACATCATCCCCTTGATACAG |
| S20L | TCGGCCATCAACAACCTGATCCCCTTGATACAG |
| S20K | TCGGCCATCAACAACAAGATCCCCTTGATACAG |
| S20M | TCGGCCATCAACAACATGATCCCCTTGATACAG |
| S20F | TCGGCCATCAACAACTTCATCCCCTTGATACAG |
| S20P | TCGGCCATCAACAACCCCATCCCCTTGATACAG |
| S20T | TCGGCCATCAACAACACCATCCCCTTGATACAG |
| S20W | TCGGCCATCAACAACTGGATCCCCTTGATACAG |
| S20Y | TCGGCCATCAACAACTACATCCCCTTGATACAG |
| S20V | TCGGCCATCAACAACGTGATCCCCTTGATACAG |
| I24A | AACAGCATCCCCTTGGCCCAGGGCAAGCTCCCG |
| I24R | AACAGCATCCCCTTGAGACAGGGCAAGCTCCCG |
| I24N | AACAGCATCCCCTTGAACCAGGGCAAGCTCCCG |
| I24D | AACAGCATCCCCTTGGACCAGGGCAAGCTCCCG |
| I24C | AACAGCATCCCCTTGTGCCAGGGCAAGCTCCCG |
| I24Q | AACAGCATCCCCTTGCAGCAGGGCAAGCTCCCG |
| I24E | AACAGCATCCCCTTGGAGCAGGGCAAGCTCCCG |
| I24G | AACAGCATCCCCTTGGGCCAGGGCAAGCTCCCG |
| I24H | AACAGCATCCCCTTGCACCAGGGCAAGCTCCCG |
| I24L | AACAGCATCCCCTTGCTGCAGGGCAAGCTCCCG |
| I24K | AACAGCATCCCCTTGAAGCAGGGCAAGCTCCCG |
| I24M | AACAGCATCCCCTTGATGCAGGGCAAGCTCCCG |
| I24F | AACAGCATCCCCTTGTTCCAGGGCAAGCTCCCG |
| I24P | AACAGCATCCCCTTGCCCCAGGGCAAGCTCCCG |
| I24S | AACAGCATCCCCTTGAGCCAGGGCAAGCTCCCG |
| I24T | AACAGCATCCCCTTGACCCAGGGCAAGCTCCCG |
| I24W | AACAGCATCCCCTTGTGGCAGGGCAAGCTCCCG |
| I24Y | AACAGCATCCCCTTGTACCAGGGCAAGCTCCCG |
| I24V | AACAGCATCCCCTTGGTGCAGGGCAAGCTCCCG |
| L28A | TTGATACAGGGCAAGGCCCCGACTCTAACCGTA |
| L28R | TTGATACAGGGCAAGAGACCGACTCTAACCGTA |
| L28N | TTGATACAGGGCAAGAACCCGACTCTAACCGTA |
| L28D | TTGATACAGGGCAAGGACCCGACTCTAACCGTA |
| L28C | TTGATACAGGGCAAGTGCCCGACTCTAACCGTA |
| L28Q | TTGATACAGGGCAAGCAGCCGACTCTAACCGTA |
| L28E | TTGATACAGGGCAAGGAGCCGACTCTAACCGTA |

| | |
|---|---|
| L28G | TTGATACAGGGCAAGGGCCCGACTCTAACCGTA |
| L28H | TTGATACAGGGCAAGCACCCGACTCTAACCGTA |
| L28I | TTGATACAGGGCAAGATCCCGACTCTAACCGTA |
| L28K | TTGATACAGGGCAAGAAGCCGACTCTAACCGTA |
| L28M | TTGATACAGGGCAAGATGCCGACTCTAACCGTA |
| L28F | TTGATACAGGGCAAGTTCCCGACTCTAACCGTA |
| L28P | TTGATACAGGGCAAGCCCCCGACTCTAACCGTA |
| L28S | TTGATACAGGGCAAGAGCCCGACTCTAACCGTA |
| L28T | TTGATACAGGGCAAGACCCCGACTCTAACCGTA |
| L28W | TTGATACAGGGCAAGTGGCCGACTCTAACCGTA |
| L28Y | TTGATACAGGGCAAGTACCCGACTCTAACCGTA |
| L28V | TTGATACAGGGCAAGGTGCCGACTCTAACCGTA |
| T32A | AAGCTCCCGACTCTAGCCGTATCTGGAAAGATC |
| T32R | AAGCTCCCGACTCTAAGAGTATCTGGAAAGATC |
| T32N | AAGCTCCCGACTCTAAACGTATCTGGAAAGATC |
| T32D | AAGCTCCCGACTCTAGACGTATCTGGAAAGATC |
| T32C | AAGCTCCCGACTCTATGCGTATCTGGAAAGATC |
| T32Q | AAGCTCCCGACTCTACAGGTATCTGGAAAGATC |
| T32E | AAGCTCCCGACTCTAGAGGTATCTGGAAAGATC |
| T32G | AAGCTCCCGACTCTAGGCGTATCTGGAAAGATC |
| T32H | AAGCTCCCGACTCTACACGTATCTGGAAAGATC |
| T32I | AAGCTCCCGACTCTAATCGTATCTGGAAAGATC |
| T32L | AAGCTCCCGACTCTACTGGTATCTGGAAAGATC |
| T32K | AAGCTCCCGACTCTAAAGGTATCTGGAAAGATC |
| T32M | AAGCTCCCGACTCTAATGGTATCTGGAAAGATC |
| T32F | AAGCTCCCGACTCTATTCGTATCTGGAAAGATC |
| T32P | AAGCTCCCGACTCTACCCGTATCTGGAAAGATC |
| T32S | AAGCTCCCGACTCTAAGCGTATCTGGAAAGATC |
| T32W | AAGCTCCCGACTCTATGGGTATCTGGAAAGATC |
| T32Y | AAGCTCCCGACTCTATACGTATCTGGAAAGATC |
| T32V | AAGCTCCCGACTCTAGTGGTATCTGGAAAGATC |
| K36A | CTAACCGTATCTGGAGCCATCCGAGTGACCGTG |
| K36R | CTAACCGTATCTGGAAGAATCCGAGTGACCGTG |
| K36N | CTAACCGTATCTGGAAACATCCGAGTGACCGTG |
| K36D | CTAACCGTATCTGGAGACATCCGAGTGACCGTG |
| K36C | CTAACCGTATCTGGATGCATCCGAGTGACCGTG |
| K36Q | CTAACCGTATCTGGACAGATCCGAGTGACCGTG |
| K36E | CTAACCGTATCTGGAGAGATCCGAGTGACCGTG |
| K36G | CTAACCGTATCTGGAGGCATCCGAGTGACCGTG |
| K36H | CTAACCGTATCTGGACACATCCGAGTGACCGTG |
| K36I | CTAACCGTATCTGGAATCATCCGAGTGACCGTG |

| | |
|---|---|
| K36L | CTAACCGTATCTGGACTGATCCGAGTGACCGTG |
| K36M | CTAACCGTATCTGGAATGATCCGAGTGACCGTG |
| K36F | CTAACCGTATCTGGATTCATCCGAGTGACCGTG |
| K36P | CTAACCGTATCTGGACCCATCCGAGTGACCGTG |
| K36S | CTAACCGTATCTGGAAGCATCCGAGTGACCGTG |
| K36T | CTAACCGTATCTGGAACCATCCGAGTGACCGTG |
| K36W | CTAACCGTATCTGGATGGATCCGAGTGACCGTG |
| K36Y | CTAACCGTATCTGGATACATCCGAGTGACCGTG |
| K36V | CTAACCGTATCTGGAGTGATCCGAGTGACCGTG |
| T40A | GGAAAGATCCGAGTGGCCGTGACTTTCTTCCTT |
| T40R | GGAAAGATCCGAGTGAGAGTGACTTTCTTCCTT |
| T40N | GGAAAGATCCGAGTGAACGTGACTTTCTTCCTT |
| T40D | GGAAAGATCCGAGTGGACGTGACTTTCTTCCTT |
| T40C | GGAAAGATCCGAGTGTGCGTGACTTTCTTCCTT |
| T40Q | GGAAAGATCCGAGTGCAGGTGACTTTCTTCCTT |
| T40E | GGAAAGATCCGAGTGGAGGTGACTTTCTTCCTT |
| T40G | GGAAAGATCCGAGTGGGCGTGACTTTCTTCCTT |
| T40H | GGAAAGATCCGAGTGCACGTGACTTTCTTCCTT |
| T40I | GGAAAGATCCGAGTGATCGTGACTTTCTTCCTT |
| T40L | GGAAAGATCCGAGTGCTGGTGACTTTCTTCCTT |
| T40K | GGAAAGATCCGAGTGAAGGTGACTTTCTTCCTT |
| T40M | GGAAAGATCCGAGTGATGGTGACTTTCTTCCTT |
| T40F | GGAAAGATCCGAGTGTTCGTGACTTTCTTCCTT |
| T40P | GGAAAGATCCGAGTGCCCGTGACTTTCTTCCTT |
| T40S | GGAAAGATCCGAGTGAGCGTGACTTTCTTCCTT |
| T40W | GGAAAGATCCGAGTGTGGGTGACTTTCTTCCTT |
| T40Y | GGAAAGATCCGAGTGTACGTGACTTTCTTCCTT |
| T40V | GGAAAGATCCGAGTGGTGGTGACTTTCTTCCTT |
| T42A | ATCCGAGTGACCGTGGCCTTCTTCCTTTTCCTA |
| T42R | ATCCGAGTGACCGTGAGATTCTTCCTTTTCCTA |
| T42N | ATCCGAGTGACCGTGAACTTCTTCCTTTTCCTA |
| T42D | ATCCGAGTGACCGTGGACTTCTTCCTTTTCCTA |
| T42C | ATCCGAGTGACCGTGTGCTTCTTCCTTTTCCTA |
| T42Q | ATCCGAGTGACCGTGCAGTTCTTCCTTTTCCTA |
| T42E | ATCCGAGTGACCGTGGAGTTCTTCCTTTTCCTA |
| T42G | ATCCGAGTGACCGTGGGCTTCTTCCTTTTCCTA |
| T42H | ATCCGAGTGACCGTGCACTTCTTCCTTTTCCTA |
| T42I | ATCCGAGTGACCGTGATCTTCTTCCTTTTCCTA |
| T42L | ATCCGAGTGACCGTGCTGTTCTTCCTTTTCCTA |
| T42K | ATCCGAGTGACCGTGAAGTTCTTCCTTTTCCTA |
| T42M | ATCCGAGTGACCGTGATGTTCTTCCTTTTCCTA |

| | |
|---|---|
| T42F | ATCCGAGTGACCGTGTTCTTCTTCCTTTTCCTA |
| T42P | ATCCGAGTGACCGTGCCCTTCTTCCTTTTCCTA |
| T42S | ATCCGAGTGACCGTGAGCTTCTTCCTTTTCCTA |
| T42W | ATCCGAGTGACCGTGTGGTTCTTCCTTTTCCTA |
| T42Y | ATCCGAGTGACCGTGTACTTCTTCCTTTTCCTA |
| T42V | ATCCGAGTGACCGTGGTGTTCTTCCTTTTCCTA |
| F44A | GTGACCGTGACTTTCGCCCTTTTCCTACTCTCT |
| F44R | GTGACCGTGACTTTCAGACTTTTCCTACTCTCT |
| F44N | GTGACCGTGACTTTCAACCTTTTCCTACTCTCT |
| F44D | GTGACCGTGACTTTCGACCTTTTCCTACTCTCT |
| F44C | GTGACCGTGACTTTCTGCCTTTTCCTACTCTCT |
| F44Q | GTGACCGTGACTTTCCAGCTTTTCCTACTCTCT |
| F44E | GTGACCGTGACTTTCGAGCTTTTCCTACTCTCT |
| F44G | GTGACCGTGACTTTCGGCCTTTTCCTACTCTCT |
| F44H | GTGACCGTGACTTTCCACCTTTTCCTACTCTCT |
| F44I | GTGACCGTGACTTTCATCCTTTTCCTACTCTCT |
| F44L | GTGACCGTGACTTTCCTGCTTTTCCTACTCTCT |
| F44K | GTGACCGTGACTTTCAAGCTTTTCCTACTCTCT |
| F44M | GTGACCGTGACTTTCATGCTTTTCCTACTCTCT |
| F44P | GTGACCGTGACTTTCCCCCTTTTCCTACTCTCT |
| F44S | GTGACCGTGACTTTCAGCCTTTTCCTACTCTCT |
| F44T | GTGACCGTGACTTTCACCCTTTTCCTACTCTCT |
| F44W | GTGACCGTGACTTTCTGGCTTTTCCTACTCTCT |
| F44Y | GTGACCGTGACTTTCTACCTTTTCCTACTCTCT |
| F44V | GTGACCGTGACTTTCGTGCTTTTCCTACTCTCT |
| S49A | TTCCTTTTCCTACTCGCCACTGCCTTCAATGCT |
| S49R | TTCCTTTTCCTACTCAGAACTGCCTTCAATGCT |
| S49N | TTCCTTTTCCTACTCAACACTGCCTTCAATGCT |
| S49D | TTCCTTTTCCTACTCGACACTGCCTTCAATGCT |
| S49C | TTCCTTTTCCTACTCTGCACTGCCTTCAATGCT |
| S49Q | TTCCTTTTCCTACTCCAGACTGCCTTCAATGCT |
| S49E | TTCCTTTTCCTACTCGAGACTGCCTTCAATGCT |
| S49G | TTCCTTTTCCTACTCGGCACTGCCTTCAATGCT |
| S49H | TTCCTTTTCCTACTCCACACTGCCTTCAATGCT |
| S49I | TTCCTTTTCCTACTCATCACTGCCTTCAATGCT |
| S49L | TTCCTTTTCCTACTCCTGACTGCCTTCAATGCT |
| S49K | TTCCTTTTCCTACTCAAGACTGCCTTCAATGCT |
| S49M | TTCCTTTTCCTACTCATGACTGCCTTCAATGCT |
| S49F | TTCCTTTTCCTACTCTTCACTGCCTTCAATGCT |
| S49P | TTCCTTTTCCTACTCCCCACTGCCTTCAATGCT |
| S49T | TTCCTTTTCCTACTCACCACTGCCTTCAATGCT |

| | |
|---|---|
| S49W | TTCCTTTTCCTACTCTGGACTGCCTTCAATGCT |
| S49Y | TTCCTTTTCCTACTCTACACTGCCTTCAATGCT |
| S49V | TTCCTTTTCCTACTCGTGACTGCCTTCAATGCT |
| A51R | TTCCTACTCTCTACTAGATTCAATGCTTCCTTC |
| A51N | TTCCTACTCTCTACTAACTTCAATGCTTCCTTC |
| A51D | TTCCTACTCTCTACTGACTTCAATGCTTCCTTC |
| A51C | TTCCTACTCTCTACTTGCTTCAATGCTTCCTTC |
| A51Q | TTCCTACTCTCTACTCAGTTCAATGCTTCCTTC |
| A51E | TTCCTACTCTCTACTGAGTTCAATGCTTCCTTC |
| A51G | TTCCTACTCTCTACTGGCTTCAATGCTTCCTTC |
| A51H | TTCCTACTCTCTACTCACTTCAATGCTTCCTTC |
| A51I | TTCCTACTCTCTACTATCTTCAATGCTTCCTTC |
| A51L | TTCCTACTCTCTACTCTGTTCAATGCTTCCTTC |
| A51K | TTCCTACTCTCTACTAAGTTCAATGCTTCCTTC |
| A51M | TTCCTACTCTCTACTATGTTCAATGCTTCCTTC |
| A51F | TTCCTACTCTCTACTTTCTTCAATGCTTCCTTC |
| A51P | TTCCTACTCTCTACTCCCTTCAATGCTTCCTTC |
| A51S | TTCCTACTCTCTACTAGCTTCAATGCTTCCTTC |
| A51T | TTCCTACTCTCTACTACCTTCAATGCTTCCTTC |
| A51W | TTCCTACTCTCTACTTGGTTCAATGCTTCCTTC |
| A51Y | TTCCTACTCTCTACTTACTTCAATGCTTCCTTC |
| A51V | TTCCTACTCTCTACTGTGTTCAATGCTTCCTTC |
| N53A | CTCTCTACTGCCTTCGCCGCTTCCTTCTTGTTG |
| N53R | CTCTCTACTGCCTTCAGAGCTTCCTTCTTGTTG |
| N53D | CTCTCTACTGCCTTCGACGCTTCCTTCTTGTTG |
| N53C | CTCTCTACTGCCTTCTGCGCTTCCTTCTTGTTG |
| N53Q | CTCTCTACTGCCTTCCAGGCTTCCTTCTTGTTG |
| N53E | CTCTCTACTGCCTTCGAGGCTTCCTTCTTGTTG |
| N53G | CTCTCTACTGCCTTCGGCGCTTCCTTCTTGTTG |
| N53H | CTCTCTACTGCCTTCCACGCTTCCTTCTTGTTG |
| N53I | CTCTCTACTGCCTTCATCGCTTCCTTCTTGTTG |
| N53L | CTCTCTACTGCCTTCCTGGCTTCCTTCTTGTTG |
| N53K | CTCTCTACTGCCTTCAAGGCTTCCTTCTTGTTG |
| N53M | CTCTCTACTGCCTTCATGGCTTCCTTCTTGTTG |
| N53F | CTCTCTACTGCCTTCTTCGCTTCCTTCTTGTTG |
| N53P | CTCTCTACTGCCTTCCCCGCTTCCTTCTTGTTG |
| N53S | CTCTCTACTGCCTTCAGCGCTTCCTTCTTGTTG |
| N53T | CTCTCTACTGCCTTCACCGCTTCCTTCTTGTTG |
| N53W | CTCTCTACTGCCTTCTGGGCTTCCTTCTTGTTG |
| N53Y | CTCTCTACTGCCTTCTACGCTTCCTTCTTGTTG |
| N53V | CTCTCTACTGCCTTCGTGGCTTCCTTCTTGTTG |

| | |
|---|---|
| T64A | AAGCTGCAGAAGTGGGCCCAGAAGAGGAAGAAA |
| T64R | AAGCTGCAGAAGTGGAGACAGAAGAGGAAGAAA |
| T64N | AAGCTGCAGAAGTGGAACCAGAAGAGGAAGAAA |
| T64D | AAGCTGCAGAAGTGGGACCAGAAGAGGAAGAAA |
| T64C | AAGCTGCAGAAGTGGTGCCAGAAGAGGAAGAAA |
| T64Q | AAGCTGCAGAAGTGGCAGCAGAAGAGGAAGAAA |
| T64E | AAGCTGCAGAAGTGGGAGCAGAAGAGGAAGAAA |
| T64G | AAGCTGCAGAAGTGGGGCCAGAAGAGGAAGAAA |
| T64H | AAGCTGCAGAAGTGGCACCAGAAGAGGAAGAAA |
| T64I | AAGCTGCAGAAGTGGATCCAGAAGAGGAAGAAA |
| T64L | AAGCTGCAGAAGTGGCTGCAGAAGAGGAAGAAA |
| T64K | AAGCTGCAGAAGTGGAAGCAGAAGAGGAAGAAA |
| T64M | AAGCTGCAGAAGTGGATGCAGAAGAGGAAGAAA |
| T64F | AAGCTGCAGAAGTGGTTCCAGAAGAGGAAGAAA |
| T64P | AAGCTGCAGAAGTGGCCCCAGAAGAGGAAGAAA |
| T64S | AAGCTGCAGAAGTGGAGCCAGAAGAGGAAGAAA |
| T64W | AAGCTGCAGAAGTGGTGGCAGAAGAGGAAGAAA |
| T64Y | AAGCTGCAGAAGTGGTACCAGAAGAGGAAGAAA |
| T64V | AAGCTGCAGAAGTGGGTGCAGAAGAGGAAGAAA |
| K68A | TGGACTCAGAAGAGGGCCAAAGGAAAAAAGCTC |
| K68R | TGGACTCAGAAGAGGAGAAAAGGAAAAAAGCTC |
| K68N | TGGACTCAGAAGAGGAACAAAGGAAAAAAGCTC |
| K68D | TGGACTCAGAAGAGGGACAAAGGAAAAAAGCTC |
| K68C | TGGACTCAGAAGAGGTGCAAAGGAAAAAAGCTC |
| K68Q | TGGACTCAGAAGAGGCAGAAAGGAAAAAAGCTC |
| K68E | TGGACTCAGAAGAGGGAGAAAGGAAAAAAGCTC |
| K68G | TGGACTCAGAAGAGGGGCAAAGGAAAAAAGCTC |
| K68H | TGGACTCAGAAGAGGCACAAAGGAAAAAAGCTC |
| K68I | TGGACTCAGAAGAGGATCAAAGGAAAAAAGCTC |
| K68L | TGGACTCAGAAGAGGCTGAAAGGAAAAAAGCTC |
| K68M | TGGACTCAGAAGAGGATGAAAGGAAAAAAGCTC |
| K68F | TGGACTCAGAAGAGGTTCAAAGGAAAAAAGCTC |
| K68P | TGGACTCAGAAGAGGCCCAAAGGAAAAAAGCTC |
| K68S | TGGACTCAGAAGAGGAGCAAAGGAAAAAAGCTC |
| K68T | TGGACTCAGAAGAGGACCAAAGGAAAAAAGCTC |
| K68W | TGGACTCAGAAGAGGTGGAAAGGAAAAAAGCTC |
| K68Y | TGGACTCAGAAGAGGTACAAAGGAAAAAAGCTC |
| K68V | TGGACTCAGAAGAGGGTGAAAGGAAAAAAGCTC |
| K72A | AGGAAGAAAGGAAAAGCCCTCTCAAGGATGAAG |
| K72R | AGGAAGAAAGGAAAAAGACTCTCAAGGATGAAG |
| K72N | AGGAAGAAAGGAAAAAACCTCTCAAGGATGAAG |

| | |
|---|---|
| K72D | AGGAAGAAAGGAAAAGACCTCTCAAGGATGAAG |
| K72C | AGGAAGAAAGGAAAATGCCTCTCAAGGATGAAG |
| K72Q | AGGAAGAAAGGAAAACAGCTCTCAAGGATGAAG |
| K72E | AGGAAGAAAGGAAAAGAGCTCTCAAGGATGAAG |
| K72G | AGGAAGAAAGGAAAAGGCCTCTCAAGGATGAAG |
| K72H | AGGAAGAAAGGAAAACACCTCTCAAGGATGAAG |
| K72I | AGGAAGAAAGGAAAAATCCTCTCAAGGATGAAG |
| K72L | AGGAAGAAAGGAAAACTGCTCTCAAGGATGAAG |
| K72M | AGGAAGAAAGGAAAAATGCTCTCAAGGATGAAG |
| K72F | AGGAAGAAAGGAAAATTCCTCTCAAGGATGAAG |
| K72P | AGGAAGAAAGGAAAACCCCTCTCAAGGATGAAG |
| K72S | AGGAAGAAAGGAAAAAGCCTCTCAAGGATGAAG |
| K72T | AGGAAGAAAGGAAAAACCCTCTCAAGGATGAAG |
| K72W | AGGAAGAAAGGAAAATGGCTCTCAAGGATGAAG |
| K72Y | AGGAAGAAAGGAAAATACCTCTCAAGGATGAAG |
| K72V | AGGAAGAAAGGAAAAGTGCTCTCAAGGATGAAG |
| M76A | AAAAAGCTCTCAAGGGCCAAGGTGCTTTTAAAG |
| M76R | AAAAAGCTCTCAAGGAGAAAGGTGCTTTTAAAG |
| M76N | AAAAAGCTCTCAAGGAACAAGGTGCTTTTAAAG |
| M76D | AAAAAGCTCTCAAGGGACAAGGTGCTTTTAAAG |
| M76C | AAAAAGCTCTCAAGGTGCAAGGTGCTTTTAAAG |
| M76Q | AAAAAGCTCTCAAGGCAGAAGGTGCTTTTAAAG |
| M76E | AAAAAGCTCTCAAGGGAGAAGGTGCTTTTAAAG |
| M76G | AAAAAGCTCTCAAGGGGCAAGGTGCTTTTAAAG |
| M76H | AAAAAGCTCTCAAGGCACAAGGTGCTTTTAAAG |
| M76I | AAAAAGCTCTCAAGGATCAAGGTGCTTTTAAAG |
| M76L | AAAAAGCTCTCAAGGCTGAAGGTGCTTTTAAAG |
| M76K | AAAAAGCTCTCAAGGAAGAAGGTGCTTTTAAAG |
| M76F | AAAAAGCTCTCAAGGTTCAAGGTGCTTTTAAAG |
| M76P | AAAAAGCTCTCAAGGCCCAAGGTGCTTTTAAAG |
| M76S | AAAAAGCTCTCAAGGAGCAAGGTGCTTTTAAAG |
| M76T | AAAAAGCTCTCAAGGACCAAGGTGCTTTTAAAG |
| M76W | AAAAAGCTCTCAAGGTGGAAGGTGCTTTTAAAG |
| M76Y | AAAAAGCTCTCAAGGTACAAGGTGCTTTTAAAG |
| M76V | AAAAAGCTCTCAAGGGTGAAGGTGCTTTTAAAG |
| T84A | CTTTTAAAGCATTTGGCCTTAGCCAACCTGCTG |
| T84R | CTTTTAAAGCATTTGAGATTAGCCAACCTGCTG |
| T84N | CTTTTAAAGCATTTGAACTTAGCCAACCTGCTG |
| T84D | CTTTTAAAGCATTTGGACTTAGCCAACCTGCTG |
| T84C | CTTTTAAAGCATTTGTGCTTAGCCAACCTGCTG |
| T84Q | CTTTTAAAGCATTTGCAGTTAGCCAACCTGCTG |

| | |
|---|---|
| T84E | CTTTTAAAGCATTTGGAGTTAGCCAACCTGCTG |
| T84G | CTTTTAAAGCATTTGGGCTTAGCCAACCTGCTG |
| T84H | CTTTTAAAGCATTTGCACTTAGCCAACCTGCTG |
| T84I | CTTTTAAAGCATTTGATCTTAGCCAACCTGCTG |
| T84L | CTTTTAAAGCATTTGCTGTTAGCCAACCTGCTG |
| T84K | CTTTTAAAGCATTTGAAGTTAGCCAACCTGCTG |
| T84M | CTTTTAAAGCATTTGATGTTAGCCAACCTGCTG |
| T84F | CTTTTAAAGCATTTGTTCTTAGCCAACCTGCTG |
| T84P | CTTTTAAAGCATTTGCCCTTAGCCAACCTGCTG |
| T84S | CTTTTAAAGCATTTGAGCTTAGCCAACCTGCTG |
| T84W | CTTTTAAAGCATTTGTGGTTAGCCAACCTGCTG |
| T84Y | CTTTTAAAGCATTTGTACTTAGCCAACCTGCTG |
| T84V | CTTTTAAAGCATTTGGTGTTAGCCAACCTGCTG |
| L85A | TTAAAGCATTTGACCGCCGCCAACCTGCTGGAG |
| L85R | TTAAAGCATTTGACCAGAGCCAACCTGCTGGAG |
| L85N | TTAAAGCATTTGACCAACGCCAACCTGCTGGAG |
| L85D | TTAAAGCATTTGACCGACGCCAACCTGCTGGAG |
| L85C | TTAAAGCATTTGACCTGCGCCAACCTGCTGGAG |
| L85Q | TTAAAGCATTTGACCCAGGCCAACCTGCTGGAG |
| L85E | TTAAAGCATTTGACCGAGGCCAACCTGCTGGAG |
| L85G | TTAAAGCATTTGACCGGCGCCAACCTGCTGGAG |
| L85H | TTAAAGCATTTGACCCACGCCAACCTGCTGGAG |
| L85I | TTAAAGCATTTGACCATCGCCAACCTGCTGGAG |
| L85K | TTAAAGCATTTGACCAAGGCCAACCTGCTGGAG |
| L85M | TTAAAGCATTTGACCATGGCCAACCTGCTGGAG |
| L85F | TTAAAGCATTTGACCTTCGCCAACCTGCTGGAG |
| L85P | TTAAAGCATTTGACCCCCGCCAACCTGCTGGAG |
| L85S | TTAAAGCATTTGACCAGCGCCAACCTGCTGGAG |
| L85T | TTAAAGCATTTGACCACCGCCAACCTGCTGGAG |
| L85W | TTAAAGCATTTGACCTGGGCCAACCTGCTGGAG |
| L85Y | TTAAAGCATTTGACCTACGCCAACCTGCTGGAG |
| L85V | TTAAAGCATTTGACCGTGGCCAACCTGCTGGAG |
| L89A | ACCTTAGCCAACCTGGCCGAGACTCTGATCGTC |
| L89R | ACCTTAGCCAACCTGAGAGAGACTCTGATCGTC |
| L89N | ACCTTAGCCAACCTGAACGAGACTCTGATCGTC |
| L89D | ACCTTAGCCAACCTGGACGAGACTCTGATCGTC |
| L89C | ACCTTAGCCAACCTGTGCGAGACTCTGATCGTC |
| L89Q | ACCTTAGCCAACCTGCAGGAGACTCTGATCGTC |
| L89E | ACCTTAGCCAACCTGGAGGAGACTCTGATCGTC |
| L89G | ACCTTAGCCAACCTGGGCGAGACTCTGATCGTC |
| L89H | ACCTTAGCCAACCTGCACGAGACTCTGATCGTC |

| | |
|---|---|
| L89I | ACCTTAGCCAACCTGATCGAGACTCTGATCGTC |
| L89K | ACCTTAGCCAACCTGAAGGAGACTCTGATCGTC |
| L89M | ACCTTAGCCAACCTGATGGAGACTCTGATCGTC |
| L89F | ACCTTAGCCAACCTGTTCGAGACTCTGATCGTC |
| L89P | ACCTTAGCCAACCTGCCCGAGACTCTGATCGTC |
| L89S | ACCTTAGCCAACCTGAGCGAGACTCTGATCGTC |
| L89T | ACCTTAGCCAACCTGACCGAGACTCTGATCGTC |
| L89W | ACCTTAGCCAACCTGTGGGAGACTCTGATCGTC |
| L89Y | ACCTTAGCCAACCTGTACGAGACTCTGATCGTC |
| L89V | ACCTTAGCCAACCTGGTGGAGACTCTGATCGTC |
| L92A | AACCTGCTGGAGACTGCCATCGTCATGCCACTG |
| L92R | AACCTGCTGGAGACTAGAATCGTCATGCCACTG |
| L92N | AACCTGCTGGAGACTAACATCGTCATGCCACTG |
| L92D | AACCTGCTGGAGACTGACATCGTCATGCCACTG |
| L92C | AACCTGCTGGAGACTTGCATCGTCATGCCACTG |
| L92Q | AACCTGCTGGAGACTCAGATCGTCATGCCACTG |
| L92E | AACCTGCTGGAGACTGAGATCGTCATGCCACTG |
| L92G | AACCTGCTGGAGACTGGCATCGTCATGCCACTG |
| L92H | AACCTGCTGGAGACTCACATCGTCATGCCACTG |
| L92I | AACCTGCTGGAGACTATCATCGTCATGCCACTG |
| L92K | AACCTGCTGGAGACTAAGATCGTCATGCCACTG |
| L92M | AACCTGCTGGAGACTATGATCGTCATGCCACTG |
| L92F | AACCTGCTGGAGACTTTCATCGTCATGCCACTG |
| L92P | AACCTGCTGGAGACTCCCATCGTCATGCCACTG |
| L92S | AACCTGCTGGAGACTAGCATCGTCATGCCACTG |
| L92T | AACCTGCTGGAGACTACCATCGTCATGCCACTG |
| L92W | AACCTGCTGGAGACTTGGATCGTCATGCCACTG |
| L92Y | AACCTGCTGGAGACTTACATCGTCATGCCACTG |
| L92V | AACCTGCTGGAGACTGTGATCGTCATGCCACTG |
| M95A | GAGACTCTGATCGTCGCCCCACTGGATGGGATG |
| M95R | GAGACTCTGATCGTCAGACCACTGGATGGGATG |
| M95N | GAGACTCTGATCGTCAACCCACTGGATGGGATG |
| M95D | GAGACTCTGATCGTCGACCCACTGGATGGGATG |
| M95C | GAGACTCTGATCGTCTGCCCACTGGATGGGATG |
| M95Q | GAGACTCTGATCGTCCAGCCACTGGATGGGATG |
| M95E | GAGACTCTGATCGTCGAGCCACTGGATGGGATG |
| M95G | GAGACTCTGATCGTCGGCCCACTGGATGGGATG |
| M95H | GAGACTCTGATCGTCCACCCACTGGATGGGATG |
| M95I | GAGACTCTGATCGTCATCCCACTGGATGGGATG |
| M95L | GAGACTCTGATCGTCCTGCCACTGGATGGGATG |
| M95K | GAGACTCTGATCGTCAAGCCACTGGATGGGATG |

| | |
|---|---|
| M95F | GAGACTCTGATCGTCTTCCCACTGGATGGGATG |
| M95P | GAGACTCTGATCGTCCCCCCACTGGATGGGATG |
| M95S | GAGACTCTGATCGTCAGCCCACTGGATGGGATG |
| M95T | GAGACTCTGATCGTCACCCCACTGGATGGGATG |
| M95W | GAGACTCTGATCGTCTGGCCACTGGATGGGATG |
| M95Y | GAGACTCTGATCGTCTACCCACTGGATGGGATG |
| M95V | GAGACTCTGATCGTCGTGCCACTGGATGGGATG |
| D98A | ATCGTCATGCCACTGGCCGGGATGTGGAATATT |
| D98R | ATCGTCATGCCACTGAGAGGGATGTGGAATATT |
| D98N | ATCGTCATGCCACTGAACGGGATGTGGAATATT |
| D98C | ATCGTCATGCCACTGTGCGGGATGTGGAATATT |
| D98Q | ATCGTCATGCCACTGCAGGGGATGTGGAATATT |
| D98E | ATCGTCATGCCACTGGAGGGGATGTGGAATATT |
| D98G | ATCGTCATGCCACTGGGCGGGATGTGGAATATT |
| D98H | ATCGTCATGCCACTGCACGGGATGTGGAATATT |
| D98I | ATCGTCATGCCACTGATCGGGATGTGGAATATT |
| D98L | ATCGTCATGCCACTGCTGGGGATGTGGAATATT |
| D98K | ATCGTCATGCCACTGAAGGGGATGTGGAATATT |
| D98M | ATCGTCATGCCACTGATGGGGATGTGGAATATT |
| D98F | ATCGTCATGCCACTGTTCGGGATGTGGAATATT |
| D98P | ATCGTCATGCCACTGCCCGGGATGTGGAATATT |
| D98S | ATCGTCATGCCACTGAGCGGGATGTGGAATATT |
| D98T | ATCGTCATGCCACTGACCGGGATGTGGAATATT |
| D98W | ATCGTCATGCCACTGTGGGGGATGTGGAATATT |
| D98Y | ATCGTCATGCCACTGTACGGGATGTGGAATATT |
| D98V | ATCGTCATGCCACTGGTGGGGATGTGGAATATT |
| V105A | ATGTGGAATATTACTGCCCAGTGGTATGCTGGG |
| V105R | ATGTGGAATATTACTAGACAGTGGTATGCTGGG |
| V105N | ATGTGGAATATTACTAACCAGTGGTATGCTGGG |
| V105D | ATGTGGAATATTACTGACCAGTGGTATGCTGGG |
| V105C | ATGTGGAATATTACTTGCCAGTGGTATGCTGGG |
| V105Q | ATGTGGAATATTACTCAGCAGTGGTATGCTGGG |
| V105E | ATGTGGAATATTACTGAGCAGTGGTATGCTGGG |
| V105G | ATGTGGAATATTACTGGCCAGTGGTATGCTGGG |
| V105H | ATGTGGAATATTACTCACCAGTGGTATGCTGGG |
| V105I | ATGTGGAATATTACTATCCAGTGGTATGCTGGG |
| V105L | ATGTGGAATATTACTCTGCAGTGGTATGCTGGG |
| V105K | ATGTGGAATATTACTAAGCAGTGGTATGCTGGG |
| V105M | ATGTGGAATATTACTATGCAGTGGTATGCTGGG |
| V105F | ATGTGGAATATTACTTTCCAGTGGTATGCTGGG |
| V105P | ATGTGGAATATTACTCCCCAGTGGTATGCTGGG |

| | |
|---|---|
| V105S | ATGTGGAATATTACTAGCCAGTGGTATGCTGGG |
| V105T | ATGTGGAATATTACTACCCAGTGGTATGCTGGG |
| V105W | ATGTGGAATATTACTTGGCAGTGGTATGCTGGG |
| V105Y | ATGTGGAATATTACTTACCAGTGGTATGCTGGG |
| Q106A | TGGAATATTACTGTTGCCTGGTATGCTGGGGAG |
| Q106R | TGGAATATTACTGTTAGATGGTATGCTGGGGAG |
| Q106N | TGGAATATTACTGTTAACTGGTATGCTGGGGAG |
| Q106D | TGGAATATTACTGTTGACTGGTATGCTGGGGAG |
| Q106C | TGGAATATTACTGTTTGCTGGTATGCTGGGGAG |
| Q106E | TGGAATATTACTGTTGAGTGGTATGCTGGGGAG |
| Q106G | TGGAATATTACTGTTGGCTGGTATGCTGGGGAG |
| Q106H | TGGAATATTACTGTTCACTGGTATGCTGGGGAG |
| Q106I | TGGAATATTACTGTTATCTGGTATGCTGGGGAG |
| Q106L | TGGAATATTACTGTTCTGTGGTATGCTGGGGAG |
| Q106K | TGGAATATTACTGTTAAGTGGTATGCTGGGGAG |
| Q106M | TGGAATATTACTGTTATGTGGTATGCTGGGGAG |
| Q106F | TGGAATATTACTGTTTTCTGGTATGCTGGGGAG |
| Q106P | TGGAATATTACTGTTCCCTGGTATGCTGGGGAG |
| Q106S | TGGAATATTACTGTTAGCTGGTATGCTGGGGAG |
| Q106T | TGGAATATTACTGTTACCTGGTATGCTGGGGAG |
| Q106W | TGGAATATTACTGTTTGGTGGTATGCTGGGGAG |
| Q106Y | TGGAATATTACTGTTTACTGGTATGCTGGGGAG |
| Q106V | TGGAATATTACTGTTGTGTGGTATGCTGGGGAG |
| Y108A | ATTACTGTTCAGTGGGCCGCTGGGGAGTTCCTC |
| Y108R | ATTACTGTTCAGTGGAGAGCTGGGGAGTTCCTC |
| Y108N | ATTACTGTTCAGTGGAACGCTGGGGAGTTCCTC |
| Y108D | ATTACTGTTCAGTGGGACGCTGGGGAGTTCCTC |
| Y108C | ATTACTGTTCAGTGGTGCGCTGGGGAGTTCCTC |
| Y108Q | ATTACTGTTCAGTGGCAGGCTGGGGAGTTCCTC |
| Y108E | ATTACTGTTCAGTGGGAGGCTGGGGAGTTCCTC |
| Y108G | ATTACTGTTCAGTGGGGCGCTGGGGAGTTCCTC |
| Y108H | ATTACTGTTCAGTGGCACGCTGGGGAGTTCCTC |
| Y108I | ATTACTGTTCAGTGGATCGCTGGGGAGTTCCTC |
| Y108L | ATTACTGTTCAGTGGCTGGCTGGGGAGTTCCTC |
| Y108K | ATTACTGTTCAGTGGAAGGCTGGGGAGTTCCTC |
| Y108M | ATTACTGTTCAGTGGATGGCTGGGGAGTTCCTC |
| Y108F | ATTACTGTTCAGTGGTTCGCTGGGGAGTTCCTC |
| Y108P | ATTACTGTTCAGTGGCCCGCTGGGGAGTTCCTC |
| Y108S | ATTACTGTTCAGTGGAGCGCTGGGGAGTTCCTC |
| Y108T | ATTACTGTTCAGTGGACCGCTGGGGAGTTCCTC |
| Y108W | ATTACTGTTCAGTGGTGGGCTGGGGAGTTCCTC |

| | |
|---|---|
| Y108V | ATTACTGTTCAGTGGGTGGCTGGGGAGTTCCTC |
| A109R | ACTGTTCAGTGGTATAGAGGGGAGTTCCTCTGC |
| A109N | ACTGTTCAGTGGTATAACGGGGAGTTCCTCTGC |
| A109D | ACTGTTCAGTGGTATGACGGGGAGTTCCTCTGC |
| A109C | ACTGTTCAGTGGTATTGCGGGGAGTTCCTCTGC |
| A109Q | ACTGTTCAGTGGTATCAGGGGGAGTTCCTCTGC |
| A109E | ACTGTTCAGTGGTATGAGGGGGAGTTCCTCTGC |
| A109G | ACTGTTCAGTGGTATGGCGGGGAGTTCCTCTGC |
| A109H | ACTGTTCAGTGGTATCACGGGGAGTTCCTCTGC |
| A109I | ACTGTTCAGTGGTATATCGGGGAGTTCCTCTGC |
| A109L | ACTGTTCAGTGGTATCTGGGGGAGTTCCTCTGC |
| A109K | ACTGTTCAGTGGTATAAGGGGGAGTTCCTCTGC |
| A109M | ACTGTTCAGTGGTATATGGGGGAGTTCCTCTGC |
| A109F | ACTGTTCAGTGGTATTTCGGGGAGTTCCTCTGC |
| A109P | ACTGTTCAGTGGTATCCCGGGGAGTTCCTCTGC |
| A109S | ACTGTTCAGTGGTATAGCGGGGAGTTCCTCTGC |
| A109T | ACTGTTCAGTGGTATACCGGGGAGTTCCTCTGC |
| A109W | ACTGTTCAGTGGTATTGGGGGGAGTTCCTCTGC |
| A109Y | ACTGTTCAGTGGTATTACGGGGAGTTCCTCTGC |
| A109V | ACTGTTCAGTGGTATGTGGGGGAGTTCCTCTGC |
| F112A | TGGTATGCTGGGGAGGCCCTCTGCAAAGTTCTC |
| F112R | TGGTATGCTGGGGAGAGACTCTGCAAAGTTCTC |
| F112N | TGGTATGCTGGGGAGAACCTCTGCAAAGTTCTC |
| F112D | TGGTATGCTGGGGAGGACCTCTGCAAAGTTCTC |
| F112C | TGGTATGCTGGGGAGTGCCTCTGCAAAGTTCTC |
| F112Q | TGGTATGCTGGGGAGCAGCTCTGCAAAGTTCTC |
| F112E | TGGTATGCTGGGGAGGAGCTCTGCAAAGTTCTC |
| F112G | TGGTATGCTGGGGAGGGCCTCTGCAAAGTTCTC |
| F112H | TGGTATGCTGGGGAGCACCTCTGCAAAGTTCTC |
| F112I | TGGTATGCTGGGGAGATCCTCTGCAAAGTTCTC |
| F112L | TGGTATGCTGGGGAGCTGCTCTGCAAAGTTCTC |
| F112K | TGGTATGCTGGGGAGAAGCTCTGCAAAGTTCTC |
| F112M | TGGTATGCTGGGGAGATGCTCTGCAAAGTTCTC |
| F112P | TGGTATGCTGGGGAGCCCCTCTGCAAAGTTCTC |
| F112S | TGGTATGCTGGGGAGAGCCTCTGCAAAGTTCTC |
| F112T | TGGTATGCTGGGGAGACCCTCTGCAAAGTTCTC |
| F112W | TGGTATGCTGGGGAGTGGCTCTGCAAAGTTCTC |
| F112Y | TGGTATGCTGGGGAGTACCTCTGCAAAGTTCTC |
| F112V | TGGTATGCTGGGGAGGTGCTCTGCAAAGTTCTC |
| V116A | GAGTTCCTCTGCAAAGCCCTCAGCTATCTGAAG |
| V116R | GAGTTCCTCTGCAAAAGACTCAGCTATCTGAAG |

| | |
|---|---|
| V116N | GAGTTCCTCTGCAAAAACCTCAGCTATCTGAAG |
| V116D | GAGTTCCTCTGCAAAGACCTCAGCTATCTGAAG |
| V116C | GAGTTCCTCTGCAAATGCCTCAGCTATCTGAAG |
| V116Q | GAGTTCCTCTGCAAACAGCTCAGCTATCTGAAG |
| V116E | GAGTTCCTCTGCAAAGAGCTCAGCTATCTGAAG |
| V116G | GAGTTCCTCTGCAAAGGCCTCAGCTATCTGAAG |
| V116H | GAGTTCCTCTGCAAACACCTCAGCTATCTGAAG |
| V116I | GAGTTCCTCTGCAAAATCCTCAGCTATCTGAAG |
| V116L | GAGTTCCTCTGCAAACTGCTCAGCTATCTGAAG |
| V116K | GAGTTCCTCTGCAAAAAGCTCAGCTATCTGAAG |
| V116M | GAGTTCCTCTGCAAAATGCTCAGCTATCTGAAG |
| V116F | GAGTTCCTCTGCAAATTCCTCAGCTATCTGAAG |
| V116P | GAGTTCCTCTGCAAACCCCTCAGCTATCTGAAG |
| V116S | GAGTTCCTCTGCAAAAGCCTCAGCTATCTGAAG |
| V116T | GAGTTCCTCTGCAAAACCCTCAGCTATCTGAAG |
| V116W | GAGTTCCTCTGCAAATGGCTCAGCTATCTGAAG |
| V116Y | GAGTTCCTCTGCAAATACCTCAGCTATCTGAAG |
| L120A | AAAGTTCTCAGCTATGCCAAGCTCTTCTCTATG |
| L120R | AAAGTTCTCAGCTATAGAAAGCTCTTCTCTATG |
| L120N | AAAGTTCTCAGCTATAACAAGCTCTTCTCTATG |
| L120D | AAAGTTCTCAGCTATGACAAGCTCTTCTCTATG |
| L120C | AAAGTTCTCAGCTATTGCAAGCTCTTCTCTATG |
| L120Q | AAAGTTCTCAGCTATCAGAAGCTCTTCTCTATG |
| L120E | AAAGTTCTCAGCTATGAGAAGCTCTTCTCTATG |
| L120G | AAAGTTCTCAGCTATGGCAAGCTCTTCTCTATG |
| L120H | AAAGTTCTCAGCTATCACAAGCTCTTCTCTATG |
| L120I | AAAGTTCTCAGCTATATCAAGCTCTTCTCTATG |
| L120K | AAAGTTCTCAGCTATAAGAAGCTCTTCTCTATG |
| L120M | AAAGTTCTCAGCTATATGAAGCTCTTCTCTATG |
| L120F | AAAGTTCTCAGCTATTTCAAGCTCTTCTCTATG |
| L120P | AAAGTTCTCAGCTATCCCAAGCTCTTCTCTATG |
| L120S | AAAGTTCTCAGCTATAGCAAGCTCTTCTCTATG |
| L120T | AAAGTTCTCAGCTATACCAAGCTCTTCTCTATG |
| L120W | AAAGTTCTCAGCTATTGGAAGCTCTTCTCTATG |
| L120Y | AAAGTTCTCAGCTATTACAAGCTCTTCTCTATG |
| L120V | AAAGTTCTCAGCTATGTGAAGCTCTTCTCTATG |
| A129R | TCTATGTATGCCCCAAGATTCATGATGGTGGTG |
| A129N | TCTATGTATGCCCCAAACTTCATGATGGTGGTG |
| A129D | TCTATGTATGCCCCAGACTTCATGATGGTGGTG |
| A129C | TCTATGTATGCCCCATGCTTCATGATGGTGGTG |
| A129Q | TCTATGTATGCCCCACAGTTCATGATGGTGGTG |

| | |
|---|---|
| A129E | TCTATGTATGCCCCAGAGTTCATGATGGTGGTG |
| A129G | TCTATGTATGCCCCAGGCTTCATGATGGTGGTG |
| A129H | TCTATGTATGCCCCACACTTCATGATGGTGGTG |
| A129I | TCTATGTATGCCCCAATCTTCATGATGGTGGTG |
| A129L | TCTATGTATGCCCCACTGTTCATGATGGTGGTG |
| A129K | TCTATGTATGCCCCAAAGTTCATGATGGTGGTG |
| A129M | TCTATGTATGCCCCAATGTTCATGATGGTGGTG |
| A129F | TCTATGTATGCCCCATTCTTCATGATGGTGGTG |
| A129P | TCTATGTATGCCCCACCCTTCATGATGGTGGTG |
| A129S | TCTATGTATGCCCCAAGCTTCATGATGGTGGTG |
| A129T | TCTATGTATGCCCCAACCTTCATGATGGTGGTG |
| A129W | TCTATGTATGCCCCATGGTTCATGATGGTGGTG |
| A129Y | TCTATGTATGCCCCATACTTCATGATGGTGGTG |
| A129V | TCTATGTATGCCCCAGTGTTCATGATGGTGGTG |
| F130A | ATGTATGCCCCAGCTGCCATGATGGTGGTGATT |
| F130R | ATGTATGCCCCAGCTAGAATGATGGTGGTGATT |
| F130N | ATGTATGCCCCAGCTAACATGATGGTGGTGATT |
| F130D | ATGTATGCCCCAGCTGACATGATGGTGGTGATT |
| F130C | ATGTATGCCCCAGCTTGCATGATGGTGGTGATT |
| F130Q | ATGTATGCCCCAGCTCAGATGATGGTGGTGATT |
| F130E | ATGTATGCCCCAGCTGAGATGATGGTGGTGATT |
| F130G | ATGTATGCCCCAGCTGGCATGATGGTGGTGATT |
| F130H | ATGTATGCCCCAGCTCACATGATGGTGGTGATT |
| F130I | ATGTATGCCCCAGCTATCATGATGGTGGTGATT |
| F130L | ATGTATGCCCCAGCTCTGATGATGGTGGTGATT |
| F130K | ATGTATGCCCCAGCTAAGATGATGGTGGTGATT |
| F130M | ATGTATGCCCCAGCTATGATGATGGTGGTGATT |
| F130P | ATGTATGCCCCAGCTCCCATGATGGTGGTGATT |
| F130S | ATGTATGCCCCAGCTAGCATGATGGTGGTGATT |
| F130T | ATGTATGCCCCAGCTACCATGATGGTGGTGATT |
| F130W | ATGTATGCCCCAGCTTGGATGATGGTGGTGATT |
| F130Y | ATGTATGCCCCAGCTTACATGATGGTGGTGATT |
| F130V | ATGTATGCCCCAGCTGTGATGATGGTGGTGATT |
| M132A | GCCCCAGCTTTCATGGCCGTGGTGATTAGCCTG |
| M132R | GCCCCAGCTTTCATGAGAGTGGTGATTAGCCTG |
| M132N | GCCCCAGCTTTCATGAACGTGGTGATTAGCCTG |
| M132D | GCCCCAGCTTTCATGGACGTGGTGATTAGCCTG |
| M132C | GCCCCAGCTTTCATGTGCGTGGTGATTAGCCTG |
| M132Q | GCCCCAGCTTTCATGCAGGTGGTGATTAGCCTG |
| M132E | GCCCCAGCTTTCATGGAGGTGGTGATTAGCCTG |
| M132G | GCCCCAGCTTTCATGGGCGTGGTGATTAGCCTG |

| | |
|---|---|
| M132H | GCCCCAGCTTTCATGCACGTGGTGATTAGCCTG |
| M132I | GCCCCAGCTTTCATGATCGTGGTGATTAGCCTG |
| M132L | GCCCCAGCTTTCATGCTGGTGGTGATTAGCCTG |
| M132K | GCCCCAGCTTTCATGAAGGTGGTGATTAGCCTG |
| M132F | GCCCCAGCTTTCATGTTCGTGGTGATTAGCCTG |
| M132P | GCCCCAGCTTTCATGCCCGTGGTGATTAGCCTG |
| M132S | GCCCCAGCTTTCATGAGCGTGGTGATTAGCCTG |
| M132T | GCCCCAGCTTTCATGACCGTGGTGATTAGCCTG |
| M132W | GCCCCAGCTTTCATGTGGGTGGTGATTAGCCTG |
| M132Y | GCCCCAGCTTTCATGTACGTGGTGATTAGCCTG |
| M132V | GCCCCAGCTTTCATGGTGGTGGTGATTAGCCTG |
| S136A | ATGATGGTGGTGATTGCCCTGGACCGCTCCCTG |
| S136R | ATGATGGTGGTGATTAGACTGGACCGCTCCCTG |
| S136N | ATGATGGTGGTGATTAACCTGGACCGCTCCCTG |
| S136D | ATGATGGTGGTGATTGACCTGGACCGCTCCCTG |
| S136C | ATGATGGTGGTGATTTGCCTGGACCGCTCCCTG |
| S136Q | ATGATGGTGGTGATTCAGCTGGACCGCTCCCTG |
| S136E | ATGATGGTGGTGATTGAGCTGGACCGCTCCCTG |
| S136G | ATGATGGTGGTGATTGGCCTGGACCGCTCCCTG |
| S136H | ATGATGGTGGTGATTCACCTGGACCGCTCCCTG |
| S136I | ATGATGGTGGTGATTATCCTGGACCGCTCCCTG |
| S136L | ATGATGGTGGTGATTCTGCTGGACCGCTCCCTG |
| S136K | ATGATGGTGGTGATTAAGCTGGACCGCTCCCTG |
| S136M | ATGATGGTGGTGATTATGCTGGACCGCTCCCTG |
| S136F | ATGATGGTGGTGATTTTCCTGGACCGCTCCCTG |
| S136P | ATGATGGTGGTGATTCCCCTGGACCGCTCCCTG |
| S136T | ATGATGGTGGTGATTACCCTGGACCGCTCCCTG |
| S136W | ATGATGGTGGTGATTTGGCTGGACCGCTCCCTG |
| S136Y | ATGATGGTGGTGATTTACCTGGACCGCTCCCTG |
| S136V | ATGATGGTGGTGATTGTGCTGGACCGCTCCCTG |
| L137A | ATGGTGGTGATTAGCGCCGACCGCTCCCTGGCC |
| L137R | ATGGTGGTGATTAGCAGAGACCGCTCCCTGGCC |
| L137N | ATGGTGGTGATTAGCAACGACCGCTCCCTGGCC |
| L137D | ATGGTGGTGATTAGCGACGACCGCTCCCTGGCC |
| L137C | ATGGTGGTGATTAGCTGCGACCGCTCCCTGGCC |
| L137Q | ATGGTGGTGATTAGCCAGGACCGCTCCCTGGCC |
| L137E | ATGGTGGTGATTAGCGAGGACCGCTCCCTGGCC |
| L137G | ATGGTGGTGATTAGCGGCGACCGCTCCCTGGCC |
| L137H | ATGGTGGTGATTAGCCACGACCGCTCCCTGGCC |
| L137I | ATGGTGGTGATTAGCATCGACCGCTCCCTGGCC |
| L137K | ATGGTGGTGATTAGCAAGGACCGCTCCCTGGCC |

| | |
|---|---|
| L137M | ATGGTGGTGATTAGCATGGACCGCTCCCTGGCC |
| L137F | ATGGTGGTGATTAGCTTCGACCGCTCCCTGGCC |
| L137P | ATGGTGGTGATTAGCCCCGACCGCTCCCTGGCC |
| L137S | ATGGTGGTGATTAGCAGCGACCGCTCCCTGGCC |
| L137T | ATGGTGGTGATTAGCACCGACCGCTCCCTGGCC |
| L137W | ATGGTGGTGATTAGCTGGGACCGCTCCCTGGCC |
| L137Y | ATGGTGGTGATTAGCTACGACCGCTCCCTGGCC |
| L137V | ATGGTGGTGATTAGCGTGGACCGCTCCCTGGCC |
| L141A | AGCCTGGACCGCTCCGCCGCCATCACTCAGCCC |
| L141R | AGCCTGGACCGCTCCAGAGCCATCACTCAGCCC |
| L141N | AGCCTGGACCGCTCCAACGCCATCACTCAGCCC |
| L141D | AGCCTGGACCGCTCCGACGCCATCACTCAGCCC |
| L141C | AGCCTGGACCGCTCCTGCGCCATCACTCAGCCC |
| L141Q | AGCCTGGACCGCTCCCAGGCCATCACTCAGCCC |
| L141E | AGCCTGGACCGCTCCGAGGCCATCACTCAGCCC |
| L141G | AGCCTGGACCGCTCCGGCGCCATCACTCAGCCC |
| L141H | AGCCTGGACCGCTCCCACGCCATCACTCAGCCC |
| L141I | AGCCTGGACCGCTCCATCGCCATCACTCAGCCC |
| L141K | AGCCTGGACCGCTCCAAGGCCATCACTCAGCCC |
| L141M | AGCCTGGACCGCTCCATGGCCATCACTCAGCCC |
| L141F | AGCCTGGACCGCTCCTTCGCCATCACTCAGCCC |
| L141P | AGCCTGGACCGCTCCCCCGCCATCACTCAGCCC |
| L141S | AGCCTGGACCGCTCCAGCGCCATCACTCAGCCC |
| L141T | AGCCTGGACCGCTCCACCGCCATCACTCAGCCC |
| L141W | AGCCTGGACCGCTCCTGGGCCATCACTCAGCCC |
| L141Y | AGCCTGGACCGCTCCTACGCCATCACTCAGCCC |
| L141V | AGCCTGGACCGCTCCGTGGCCATCACTCAGCCC |
| A148R | ATCACTCAGCCCCTTAGAGTACAAAGCAACAGC |
| A148N | ATCACTCAGCCCCTTAACGTACAAAGCAACAGC |
| A148D | ATCACTCAGCCCCTTGACGTACAAAGCAACAGC |
| A148C | ATCACTCAGCCCCTTTGCGTACAAAGCAACAGC |
| A148Q | ATCACTCAGCCCCTTCAGGTACAAAGCAACAGC |
| A148E | ATCACTCAGCCCCTTGAGGTACAAAGCAACAGC |
| A148G | ATCACTCAGCCCCTTGGCGTACAAAGCAACAGC |
| A148H | ATCACTCAGCCCCTTCACGTACAAAGCAACAGC |
| A148I | ATCACTCAGCCCCTTATCGTACAAAGCAACAGC |
| A148L | ATCACTCAGCCCCTTCTGGTACAAAGCAACAGC |
| A148K | ATCACTCAGCCCCTTAAGGTACAAAGCAACAGC |
| A148M | ATCACTCAGCCCCTTATGGTACAAAGCAACAGC |
| A148F | ATCACTCAGCCCCTTTTCGTACAAAGCAACAGC |
| A148P | ATCACTCAGCCCCTTCCCGTACAAAGCAACAGC |

| | |
|---|---|
| A148S | ATCACTCAGCCCCTTAGCGTACAAAGCAACAGC |
| A148T | ATCACTCAGCCCCTTACCGTACAAAGCAACAGC |
| A148W | ATCACTCAGCCCCTTTGGGTACAAAGCAACAGC |
| A148Y | ATCACTCAGCCCCTTTACGTACAAAGCAACAGC |
| A148V | ATCACTCAGCCCCTTGTGGTACAAAGCAACAGC |
| N152A | CTTGCTGTACAAAGCGCCAGCAAGCTTGAACAG |
| N152R | CTTGCTGTACAAAGCAGAAGCAAGCTTGAACAG |
| N152D | CTTGCTGTACAAAGCGACAGCAAGCTTGAACAG |
| N152C | CTTGCTGTACAAAGCTGCAGCAAGCTTGAACAG |
| N152Q | CTTGCTGTACAAAGCCAGAGCAAGCTTGAACAG |
| N152E | CTTGCTGTACAAAGCGAGAGCAAGCTTGAACAG |
| N152G | CTTGCTGTACAAAGCGGCAGCAAGCTTGAACAG |
| N152H | CTTGCTGTACAAAGCCACAGCAAGCTTGAACAG |
| N152I | CTTGCTGTACAAAGCATCAGCAAGCTTGAACAG |
| N152L | CTTGCTGTACAAAGCCTGAGCAAGCTTGAACAG |
| N152K | CTTGCTGTACAAAGCAAGAGCAAGCTTGAACAG |
| N152M | CTTGCTGTACAAAGCATGAGCAAGCTTGAACAG |
| N152F | CTTGCTGTACAAAGCTTCAGCAAGCTTGAACAG |
| N152P | CTTGCTGTACAAAGCCCCAGCAAGCTTGAACAG |
| N152S | CTTGCTGTACAAAGCAGCAGCAAGCTTGAACAG |
| N152T | CTTGCTGTACAAAGCACCAGCAAGCTTGAACAG |
| N152W | CTTGCTGTACAAAGCTGGAGCAAGCTTGAACAG |
| N152Y | CTTGCTGTACAAAGCTACAGCAAGCTTGAACAG |
| N152V | CTTGCTGTACAAAGCGTGAGCAAGCTTGAACAG |
| E156A | AGCAACAGCAAGCTTGCCCAGTCTATGATCAGC |
| E156R | AGCAACAGCAAGCTTAGACAGTCTATGATCAGC |
| E156N | AGCAACAGCAAGCTTAACCAGTCTATGATCAGC |
| E156D | AGCAACAGCAAGCTTGACCAGTCTATGATCAGC |
| E156C | AGCAACAGCAAGCTTTGCCAGTCTATGATCAGC |
| E156Q | AGCAACAGCAAGCTTCAGCAGTCTATGATCAGC |
| E156G | AGCAACAGCAAGCTTGGCCAGTCTATGATCAGC |
| E156H | AGCAACAGCAAGCTTCACCAGTCTATGATCAGC |
| E156I | AGCAACAGCAAGCTTATCCAGTCTATGATCAGC |
| E156L | AGCAACAGCAAGCTTCTGCAGTCTATGATCAGC |
| E156K | AGCAACAGCAAGCTTAAGCAGTCTATGATCAGC |
| E156M | AGCAACAGCAAGCTTATGCAGTCTATGATCAGC |
| E156F | AGCAACAGCAAGCTTTTCCAGTCTATGATCAGC |
| E156P | AGCAACAGCAAGCTTCCCCAGTCTATGATCAGC |
| E156S | AGCAACAGCAAGCTTAGCCAGTCTATGATCAGC |
| E156T | AGCAACAGCAAGCTTACCCAGTCTATGATCAGC |
| E156W | AGCAACAGCAAGCTTTGGCAGTCTATGATCAGC |

| | |
|---|---|
| E156Y | AGCAACAGCAAGCTTTACCAGTCTATGATCAGC |
| E156V | AGCAACAGCAAGCTTGTGCAGTCTATGATCAGC |
| I160A | CTTGAACAGTCTATGGCCAGCCTGGCCTGGATT |
| I160R | CTTGAACAGTCTATGAGAAGCCTGGCCTGGATT |
| I160N | CTTGAACAGTCTATGAACAGCCTGGCCTGGATT |
| I160D | CTTGAACAGTCTATGGACAGCCTGGCCTGGATT |
| I160C | CTTGAACAGTCTATGTGCAGCCTGGCCTGGATT |
| I160Q | CTTGAACAGTCTATGCAGAGCCTGGCCTGGATT |
| I160E | CTTGAACAGTCTATGGAGAGCCTGGCCTGGATT |
| I160G | CTTGAACAGTCTATGGGCAGCCTGGCCTGGATT |
| I160H | CTTGAACAGTCTATGCACAGCCTGGCCTGGATT |
| I160L | CTTGAACAGTCTATGCTGAGCCTGGCCTGGATT |
| I160K | CTTGAACAGTCTATGAAGAGCCTGGCCTGGATT |
| I160M | CTTGAACAGTCTATGATGAGCCTGGCCTGGATT |
| I160F | CTTGAACAGTCTATGTTCAGCCTGGCCTGGATT |
| I160P | CTTGAACAGTCTATGCCCAGCCTGGCCTGGATT |
| I160S | CTTGAACAGTCTATGAGCAGCCTGGCCTGGATT |
| I160T | CTTGAACAGTCTATGACCAGCCTGGCCTGGATT |
| I160W | CTTGAACAGTCTATGTGGAGCCTGGCCTGGATT |
| I160Y | CTTGAACAGTCTATGTACAGCCTGGCCTGGATT |
| I160V | CTTGAACAGTCTATGGTGAGCCTGGCCTGGATT |
| A163R | TCTATGATCAGCCTGAGATGGATTCTCAGCATT |
| A163N | TCTATGATCAGCCTGAACTGGATTCTCAGCATT |
| A163D | TCTATGATCAGCCTGGACTGGATTCTCAGCATT |
| A163C | TCTATGATCAGCCTGTGCTGGATTCTCAGCATT |
| A163Q | TCTATGATCAGCCTGCAGTGGATTCTCAGCATT |
| A163E | TCTATGATCAGCCTGGAGTGGATTCTCAGCATT |
| A163G | TCTATGATCAGCCTGGGCTGGATTCTCAGCATT |
| A163H | TCTATGATCAGCCTGCACTGGATTCTCAGCATT |
| A163I | TCTATGATCAGCCTGATCTGGATTCTCAGCATT |
| A163L | TCTATGATCAGCCTGCTGTGGATTCTCAGCATT |
| A163K | TCTATGATCAGCCTGAAGTGGATTCTCAGCATT |
| A163M | TCTATGATCAGCCTGATGTGGATTCTCAGCATT |
| A163F | TCTATGATCAGCCTGTTCTGGATTCTCAGCATT |
| A163P | TCTATGATCAGCCTGCCCTGGATTCTCAGCATT |
| A163S | TCTATGATCAGCCTGAGCTGGATTCTCAGCATT |
| A163T | TCTATGATCAGCCTGACCTGGATTCTCAGCATT |
| A163W | TCTATGATCAGCCTGTGGTGGATTCTCAGCATT |
| A163Y | TCTATGATCAGCCTGTACTGGATTCTCAGCATT |
| A163V | TCTATGATCAGCCTGGTGTGGATTCTCAGCATT |
| I165A | ATCAGCCTGGCCTGGGCCCTCAGCATTGTCTTT |

| | |
|---|---|
| I165R | ATCAGCCTGGCCTGGAGACTCAGCATTGTCTTT |
| I165N | ATCAGCCTGGCCTGGAACCTCAGCATTGTCTTT |
| I165D | ATCAGCCTGGCCTGGGACCTCAGCATTGTCTTT |
| I165C | ATCAGCCTGGCCTGGTGCCTCAGCATTGTCTTT |
| I165Q | ATCAGCCTGGCCTGGCAGCTCAGCATTGTCTTT |
| I165E | ATCAGCCTGGCCTGGGAGCTCAGCATTGTCTTT |
| I165G | ATCAGCCTGGCCTGGGGCCTCAGCATTGTCTTT |
| I165H | ATCAGCCTGGCCTGGCACCTCAGCATTGTCTTT |
| I165L | ATCAGCCTGGCCTGGCTGCTCAGCATTGTCTTT |
| I165K | ATCAGCCTGGCCTGGAAGCTCAGCATTGTCTTT |
| I165M | ATCAGCCTGGCCTGGATGCTCAGCATTGTCTTT |
| I165F | ATCAGCCTGGCCTGGTTCCTCAGCATTGTCTTT |
| I165P | ATCAGCCTGGCCTGGCCCCTCAGCATTGTCTTT |
| I165S | ATCAGCCTGGCCTGGAGCCTCAGCATTGTCTTT |
| I165T | ATCAGCCTGGCCTGGACCCTCAGCATTGTCTTT |
| I165W | ATCAGCCTGGCCTGGTGGCTCAGCATTGTCTTT |
| I165Y | ATCAGCCTGGCCTGGTACCTCAGCATTGTCTTT |
| I165V | ATCAGCCTGGCCTGGGTGCTCAGCATTGTCTTT |
| I168A | GCCTGGATTCTCAGCGCCGTCTTTGCAGGACCA |
| I168R | GCCTGGATTCTCAGCAGAGTCTTTGCAGGACCA |
| I168N | GCCTGGATTCTCAGCAACGTCTTTGCAGGACCA |
| I168D | GCCTGGATTCTCAGCGACGTCTTTGCAGGACCA |
| I168C | GCCTGGATTCTCAGCTGCGTCTTTGCAGGACCA |
| I168Q | GCCTGGATTCTCAGCCAGGTCTTTGCAGGACCA |
| I168E | GCCTGGATTCTCAGCGAGGTCTTTGCAGGACCA |
| I168G | GCCTGGATTCTCAGCGGCGTCTTTGCAGGACCA |
| I168H | GCCTGGATTCTCAGCCACGTCTTTGCAGGACCA |
| I168L | GCCTGGATTCTCAGCCTGGTCTTTGCAGGACCA |
| I168K | GCCTGGATTCTCAGCAAGGTCTTTGCAGGACCA |
| I168M | GCCTGGATTCTCAGCATGGTCTTTGCAGGACCA |
| I168F | GCCTGGATTCTCAGCTTCGTCTTTGCAGGACCA |
| I168P | GCCTGGATTCTCAGCCCCGTCTTTGCAGGACCA |
| I168S | GCCTGGATTCTCAGCAGCGTCTTTGCAGGACCA |
| I168T | GCCTGGATTCTCAGCACCGTCTTTGCAGGACCA |
| I168W | GCCTGGATTCTCAGCTGGGTCTTTGCAGGACCA |
| I168Y | GCCTGGATTCTCAGCTACGTCTTTGCAGGACCA |
| I168V | GCCTGGATTCTCAGCGTGGTCTTTGCAGGACCA |
| Q174A | GTCTTTGCAGGACCAGCCTTATATATCTTCAGG |
| Q174R | GTCTTTGCAGGACCAAGATTATATATCTTCAGG |
| Q174N | GTCTTTGCAGGACCAAACTTATATATCTTCAGG |
| Q174D | GTCTTTGCAGGACCAGACTTATATATCTTCAGG |

| | |
|---|---|
| Q174C | GTCTTTGCAGGACCATGCTTATATATCTTCAGG |
| Q174E | GTCTTTGCAGGACCAGAGTTATATATCTTCAGG |
| Q174G | GTCTTTGCAGGACCAGGCTTATATATCTTCAGG |
| Q174H | GTCTTTGCAGGACCACACTTATATATCTTCAGG |
| Q174I | GTCTTTGCAGGACCAATCTTATATATCTTCAGG |
| Q174L | GTCTTTGCAGGACCACTGTTATATATCTTCAGG |
| Q174K | GTCTTTGCAGGACCAAAGTTATATATCTTCAGG |
| Q174M | GTCTTTGCAGGACCAATGTTATATATCTTCAGG |
| Q174F | GTCTTTGCAGGACCATTCTTATATATCTTCAGG |
| Q174P | GTCTTTGCAGGACCACCCTTATATATCTTCAGG |
| Q174S | GTCTTTGCAGGACCAAGCTTATATATCTTCAGG |
| Q174T | GTCTTTGCAGGACCAACCTTATATATCTTCAGG |
| Q174W | GTCTTTGCAGGACCATGGTTATATATCTTCAGG |
| Q174Y | GTCTTTGCAGGACCATACTTATATATCTTCAGG |
| Q174V | GTCTTTGCAGGACCAGTGTTATATATCTTCAGG |
| L175A | TTTGCAGGACCACAGGCCTATATCTTCAGGATG |
| L175R | TTTGCAGGACCACAGAGATATATCTTCAGGATG |
| L175N | TTTGCAGGACCACAGAACTATATCTTCAGGATG |
| L175D | TTTGCAGGACCACAGGACTATATCTTCAGGATG |
| L175C | TTTGCAGGACCACAGTGCTATATCTTCAGGATG |
| L175Q | TTTGCAGGACCACAGCAGTATATCTTCAGGATG |
| L175E | TTTGCAGGACCACAGGAGTATATCTTCAGGATG |
| L175G | TTTGCAGGACCACAGGGCTATATCTTCAGGATG |
| L175H | TTTGCAGGACCACAGCACTATATCTTCAGGATG |
| L175I | TTTGCAGGACCACAGATCTATATCTTCAGGATG |
| L175K | TTTGCAGGACCACAGAAGTATATCTTCAGGATG |
| L175M | TTTGCAGGACCACAGATGTATATCTTCAGGATG |
| L175F | TTTGCAGGACCACAGTTCTATATCTTCAGGATG |
| L175P | TTTGCAGGACCACAGCCCTATATCTTCAGGATG |
| L175S | TTTGCAGGACCACAGAGCTATATCTTCAGGATG |
| L175T | TTTGCAGGACCACAGACCTATATCTTCAGGATG |
| L175W | TTTGCAGGACCACAGTGGTATATCTTCAGGATG |
| L175Y | TTTGCAGGACCACAGTACTATATCTTCAGGATG |
| L175V | TTTGCAGGACCACAGGTGTATATCTTCAGGATG |
| I177A | GGACCACAGTTATATGCCTTCAGGATGATCTAC |
| I177R | GGACCACAGTTATATAGATTCAGGATGATCTAC |
| I177N | GGACCACAGTTATATAACTTCAGGATGATCTAC |
| I177D | GGACCACAGTTATATGACTTCAGGATGATCTAC |
| I177C | GGACCACAGTTATATTGCTTCAGGATGATCTAC |
| I177Q | GGACCACAGTTATATCAGTTCAGGATGATCTAC |
| I177E | GGACCACAGTTATATGAGTTCAGGATGATCTAC |

| | |
|---|---|
| I177G | GGACCACAGTTATATGGCTTCAGGATGATCTAC |
| I177H | GGACCACAGTTATATCACTTCAGGATGATCTAC |
| I177L | GGACCACAGTTATATCTGTTCAGGATGATCTAC |
| I177K | GGACCACAGTTATATAAGTTCAGGATGATCTAC |
| I177M | GGACCACAGTTATATATGTTCAGGATGATCTAC |
| I177F | GGACCACAGTTATATTTCTTCAGGATGATCTAC |
| I177P | GGACCACAGTTATATCCCTTCAGGATGATCTAC |
| I177S | GGACCACAGTTATATAGCTTCAGGATGATCTAC |
| I177T | GGACCACAGTTATATACCTTCAGGATGATCTAC |
| I177W | GGACCACAGTTATATTGGTTCAGGATGATCTAC |
| I177Y | GGACCACAGTTATATTACTTCAGGATGATCTAC |
| I177V | GGACCACAGTTATATGTGTTCAGGATGATCTAC |
| G188A | CTAGCAGACGGCTCTGCCCCCACAGTCTTCTCG |
| G188R | CTAGCAGACGGCTCTAGACCCACAGTCTTCTCG |
| G188N | CTAGCAGACGGCTCTAACCCCACAGTCTTCTCG |
| G188D | CTAGCAGACGGCTCTGACCCCACAGTCTTCTCG |
| G188C | CTAGCAGACGGCTCTTGCCCCACAGTCTTCTCG |
| G188Q | CTAGCAGACGGCTCTCAGCCCACAGTCTTCTCG |
| G188E | CTAGCAGACGGCTCTGAGCCCACAGTCTTCTCG |
| G188H | CTAGCAGACGGCTCTCACCCCACAGTCTTCTCG |
| G188I | CTAGCAGACGGCTCTATCCCCACAGTCTTCTCG |
| G188L | CTAGCAGACGGCTCTCTGCCCACAGTCTTCTCG |
| G188K | CTAGCAGACGGCTCTAAGCCCACAGTCTTCTCG |
| G188M | CTAGCAGACGGCTCTATGCCCACAGTCTTCTCG |
| G188F | CTAGCAGACGGCTCTTTCCCCACAGTCTTCTCG |
| G188P | CTAGCAGACGGCTCTCCCCCCACAGTCTTCTCG |
| G188S | CTAGCAGACGGCTCTAGCCCCACAGTCTTCTCG |
| G188T | CTAGCAGACGGCTCTACCCCCACAGTCTTCTCG |
| G188W | CTAGCAGACGGCTCTTGGCCCACAGTCTTCTCG |
| G188Y | CTAGCAGACGGCTCTTACCCCACAGTCTTCTCG |
| G188V | CTAGCAGACGGCTCTGTGCCCACAGTCTTCTCG |
| F192A | TCTGGGCCCACAGTCGCCTCGCAATGTGTGACC |
| F192R | TCTGGGCCCACAGTCAGATCGCAATGTGTGACC |
| F192N | TCTGGGCCCACAGTCAACTCGCAATGTGTGACC |
| F192D | TCTGGGCCCACAGTCGACTCGCAATGTGTGACC |
| F192C | TCTGGGCCCACAGTCTGCTCGCAATGTGTGACC |
| F192Q | TCTGGGCCCACAGTCCAGTCGCAATGTGTGACC |
| F192E | TCTGGGCCCACAGTCGAGTCGCAATGTGTGACC |
| F192G | TCTGGGCCCACAGTCGGCTCGCAATGTGTGACC |
| F192H | TCTGGGCCCACAGTCCACTCGCAATGTGTGACC |
| F192I | TCTGGGCCCACAGTCATCTCGCAATGTGTGACC |

| | |
|---|---|
| F192L | TCTGGGCCCACAGTCCTGTCGCAATGTGTGACC |
| F192K | TCTGGGCCCACAGTCAAGTCGCAATGTGTGACC |
| F192M | TCTGGGCCCACAGTCATGTCGCAATGTGTGACC |
| F192P | TCTGGGCCCACAGTCCCCTCGCAATGTGTGACC |
| F192S | TCTGGGCCCACAGTCAGCTCGCAATGTGTGACC |
| F192T | TCTGGGCCCACAGTCACCTCGCAATGTGTGACC |
| F192W | TCTGGGCCCACAGTCTGGTCGCAATGTGTGACC |
| F192Y | TCTGGGCCCACAGTCTACTCGCAATGTGTGACC |
| F192V | TCTGGGCCCACAGTCGTGTCGCAATGTGTGACC |
| V196A | GTCTTCTCGCAATGTGCCACCCACTGCAGCTTT |
| V196R | GTCTTCTCGCAATGTAGAACCCACTGCAGCTTT |
| V196N | GTCTTCTCGCAATGTAACACCCACTGCAGCTTT |
| V196D | GTCTTCTCGCAATGTGACACCCACTGCAGCTTT |
| V196C | GTCTTCTCGCAATGTTGCACCCACTGCAGCTTT |
| V196Q | GTCTTCTCGCAATGTCAGACCCACTGCAGCTTT |
| V196E | GTCTTCTCGCAATGTGAGACCCACTGCAGCTTT |
| V196G | GTCTTCTCGCAATGTGGCACCCACTGCAGCTTT |
| V196H | GTCTTCTCGCAATGTCACACCCACTGCAGCTTT |
| V196I | GTCTTCTCGCAATGTATCACCCACTGCAGCTTT |
| V196L | GTCTTCTCGCAATGTCTGACCCACTGCAGCTTT |
| V196K | GTCTTCTCGCAATGTAAGACCCACTGCAGCTTT |
| V196M | GTCTTCTCGCAATGTATGACCCACTGCAGCTTT |
| V196F | GTCTTCTCGCAATGTTTCACCCACTGCAGCTTT |
| V196P | GTCTTCTCGCAATGTCCCACCCACTGCAGCTTT |
| V196S | GTCTTCTCGCAATGTAGCACCCACTGCAGCTTT |
| V196T | GTCTTCTCGCAATGTACCACCCACTGCAGCTTT |
| V196W | GTCTTCTCGCAATGTTGGACCCACTGCAGCTTT |
| V196Y | GTCTTCTCGCAATGTTACACCCACTGCAGCTTT |
| S200A | TGTGTGACCCACTGCGCCTTTCCACAGTGGTGG |
| S200R | TGTGTGACCCACTGCAGATTTCCACAGTGGTGG |
| S200N | TGTGTGACCCACTGCAACTTTCCACAGTGGTGG |
| S200D | TGTGTGACCCACTGCGACTTTCCACAGTGGTGG |
| S200C | TGTGTGACCCACTGCTGCTTTCCACAGTGGTGG |
| S200Q | TGTGTGACCCACTGCCAGTTTCCACAGTGGTGG |
| S200E | TGTGTGACCCACTGCGAGTTTCCACAGTGGTGG |
| S200G | TGTGTGACCCACTGCGGCTTTCCACAGTGGTGG |
| S200H | TGTGTGACCCACTGCCACTTTCCACAGTGGTGG |
| S200I | TGTGTGACCCACTGCATCTTTCCACAGTGGTGG |
| S200L | TGTGTGACCCACTGCCTGTTTCCACAGTGGTGG |
| S200K | TGTGTGACCCACTGCAAGTTTCCACAGTGGTGG |
| S200M | TGTGTGACCCACTGCATGTTTCCACAGTGGTGG |

| | |
|---|---|
| S200F | TGTGTGACCCACTGCTTCTTTCCACAGTGGTGG |
| S200P | TGTGTGACCCACTGCCCCTTTCCACAGTGGTGG |
| S200T | TGTGTGACCCACTGCACCTTTCCACAGTGGTGG |
| S200W | TGTGTGACCCACTGCTGGTTTCCACAGTGGTGG |
| S200Y | TGTGTGACCCACTGCTACTTTCCACAGTGGTGG |
| S200V | TGTGTGACCCACTGCGTGTTTCCACAGTGGTGG |
| W204A | TGCAGCTTTCCACAGGCCTGGCATCAGGCCTTC |
| W204R | TGCAGCTTTCCACAGAGATGGCATCAGGCCTTC |
| W204N | TGCAGCTTTCCACAGAACTGGCATCAGGCCTTC |
| W204D | TGCAGCTTTCCACAGGACTGGCATCAGGCCTTC |
| W204C | TGCAGCTTTCCACAGTGCTGGCATCAGGCCTTC |
| W204Q | TGCAGCTTTCCACAGCAGTGGCATCAGGCCTTC |
| W204E | TGCAGCTTTCCACAGGAGTGGCATCAGGCCTTC |
| W204G | TGCAGCTTTCCACAGGGCTGGCATCAGGCCTTC |
| W204H | TGCAGCTTTCCACAGCACTGGCATCAGGCCTTC |
| W204I | TGCAGCTTTCCACAGATCTGGCATCAGGCCTTC |
| W204L | TGCAGCTTTCCACAGCTGTGGCATCAGGCCTTC |
| W204K | TGCAGCTTTCCACAGAAGTGGCATCAGGCCTTC |
| W204M | TGCAGCTTTCCACAGATGTGGCATCAGGCCTTC |
| W204F | TGCAGCTTTCCACAGTTCTGGCATCAGGCCTTC |
| W204P | TGCAGCTTTCCACAGCCCTGGCATCAGGCCTTC |
| W204S | TGCAGCTTTCCACAGAGCTGGCATCAGGCCTTC |
| W204T | TGCAGCTTTCCACAGACCTGGCATCAGGCCTTC |
| W204Y | TGCAGCTTTCCACAGTACTGGCATCAGGCCTTC |
| W204V | TGCAGCTTTCCACAGGTGTGGCATCAGGCCTTC |
| A208R | CAGTGGTGGCATCAGAGATTCTACAACTTTTTC |
| A208N | CAGTGGTGGCATCAGAACTTCTACAACTTTTTC |
| A208D | CAGTGGTGGCATCAGGACTTCTACAACTTTTTC |
| A208C | CAGTGGTGGCATCAGTGCTTCTACAACTTTTTC |
| A208Q | CAGTGGTGGCATCAGCAGTTCTACAACTTTTTC |
| A208E | CAGTGGTGGCATCAGGAGTTCTACAACTTTTTC |
| A208G | CAGTGGTGGCATCAGGGCTTCTACAACTTTTTC |
| A208H | CAGTGGTGGCATCAGCACTTCTACAACTTTTTC |
| A208I | CAGTGGTGGCATCAGATCTTCTACAACTTTTTC |
| A208L | CAGTGGTGGCATCAGCTGTTCTACAACTTTTTC |
| A208K | CAGTGGTGGCATCAGAAGTTCTACAACTTTTTC |
| A208M | CAGTGGTGGCATCAGATGTTCTACAACTTTTTC |
| A208F | CAGTGGTGGCATCAGTTCTTCTACAACTTTTTC |
| A208P | CAGTGGTGGCATCAGCCCTTCTACAACTTTTTC |
| A208S | CAGTGGTGGCATCAGAGCTTCTACAACTTTTTC |
| A208T | CAGTGGTGGCATCAGACCTTCTACAACTTTTTC |

| | |
|---|---|
| A208W | CAGTGGTGGCATCAGTGGTTCTACAACTTTTTC |
| A208Y | CAGTGGTGGCATCAGTACTTCTACAACTTTTTC |
| A208V | CAGTGGTGGCATCAGGTGTTCTACAACTTTTTC |
| F219A | ACCTTCGGCTGCCTCGCCATCATCCCCCTCCTC |
| F219R | ACCTTCGGCTGCCTCAGAATCATCCCCCTCCTC |
| F219N | ACCTTCGGCTGCCTCAACATCATCCCCCTCCTC |
| F219D | ACCTTCGGCTGCCTCGACATCATCCCCCTCCTC |
| F219C | ACCTTCGGCTGCCTCTGCATCATCCCCCTCCTC |
| F219Q | ACCTTCGGCTGCCTCCAGATCATCCCCCTCCTC |
| F219E | ACCTTCGGCTGCCTCGAGATCATCCCCCTCCTC |
| F219G | ACCTTCGGCTGCCTCGGCATCATCCCCCTCCTC |
| F219H | ACCTTCGGCTGCCTCCACATCATCCCCCTCCTC |
| F219I | ACCTTCGGCTGCCTCATCATCATCCCCCTCCTC |
| F219L | ACCTTCGGCTGCCTCCTGATCATCCCCCTCCTC |
| F219K | ACCTTCGGCTGCCTCAAGATCATCCCCCTCCTC |
| F219M | ACCTTCGGCTGCCTCATGATCATCCCCCTCCTC |
| F219P | ACCTTCGGCTGCCTCCCCATCATCCCCCTCCTC |
| F219S | ACCTTCGGCTGCCTCAGCATCATCCCCCTCCTC |
| F219T | ACCTTCGGCTGCCTCACCATCATCCCCCTCCTC |
| F219W | ACCTTCGGCTGCCTCTGGATCATCCCCCTCCTC |
| F219Y | ACCTTCGGCTGCCTCTACATCATCCCCCTCCTC |
| F219V | ACCTTCGGCTGCCTCGTGATCATCCCCCTCCTC |
| I221A | GGCTGCCTCTTCATCGCCCCCCTCCTCATCATG |
| I221R | GGCTGCCTCTTCATCAGACCCCTCCTCATCATG |
| I221N | GGCTGCCTCTTCATCAACCCCCTCCTCATCATG |
| I221D | GGCTGCCTCTTCATCGACCCCCTCCTCATCATG |
| I221C | GGCTGCCTCTTCATCTGCCCCCTCCTCATCATG |
| I221Q | GGCTGCCTCTTCATCCAGCCCCTCCTCATCATG |
| I221E | GGCTGCCTCTTCATCGAGCCCCTCCTCATCATG |
| I221G | GGCTGCCTCTTCATCGGCCCCCTCCTCATCATG |
| I221H | GGCTGCCTCTTCATCCACCCCCTCCTCATCATG |
| I221L | GGCTGCCTCTTCATCCTGCCCCTCCTCATCATG |
| I221K | GGCTGCCTCTTCATCAAGCCCCTCCTCATCATG |
| I221M | GGCTGCCTCTTCATCATGCCCCTCCTCATCATG |
| I221F | GGCTGCCTCTTCATCTTCCCCCTCCTCATCATG |
| I221P | GGCTGCCTCTTCATCCCCCCCCTCCTCATCATG |
| I221S | GGCTGCCTCTTCATCAGCCCCCTCCTCATCATG |
| I221T | GGCTGCCTCTTCATCACCCCCCTCCTCATCATG |
| I221W | GGCTGCCTCTTCATCTGGCCCCTCCTCATCATG |
| I221Y | GGCTGCCTCTTCATCTACCCCCTCCTCATCATG |
| I221V | GGCTGCCTCTTCATCGTGCCCCTCCTCATCATG |

| | |
|---|---|
| M226A | ATCCCCCTCCTCATCGCCCTAATCTGCAATGCC |
| M226R | ATCCCCCTCCTCATCAGACTAATCTGCAATGCC |
| M226N | ATCCCCCTCCTCATCAACCTAATCTGCAATGCC |
| M226D | ATCCCCCTCCTCATCGACCTAATCTGCAATGCC |
| M226C | ATCCCCCTCCTCATCTGCCTAATCTGCAATGCC |
| M226Q | ATCCCCCTCCTCATCCAGCTAATCTGCAATGCC |
| M226E | ATCCCCCTCCTCATCGAGCTAATCTGCAATGCC |
| M226G | ATCCCCCTCCTCATCGGCCTAATCTGCAATGCC |
| M226H | ATCCCCCTCCTCATCCACCTAATCTGCAATGCC |
| M226I | ATCCCCCTCCTCATCATCCTAATCTGCAATGCC |
| M226L | ATCCCCCTCCTCATCCTGCTAATCTGCAATGCC |
| M226K | ATCCCCCTCCTCATCAAGCTAATCTGCAATGCC |
| M226F | ATCCCCCTCCTCATCTTCCTAATCTGCAATGCC |
| M226P | ATCCCCCTCCTCATCCCCCTAATCTGCAATGCC |
| M226S | ATCCCCCTCCTCATCAGCCTAATCTGCAATGCC |
| M226T | ATCCCCCTCCTCATCACCCTAATCTGCAATGCC |
| M226W | ATCCCCCTCCTCATCTGGCTAATCTGCAATGCC |
| M226Y | ATCCCCCTCCTCATCTACCTAATCTGCAATGCC |
| M226V | ATCCCCCTCCTCATCGTGCTAATCTGCAATGCC |
| I228A | CTCCTCATCATGCTAGCCTGCAATGCCAAAATC |
| I228R | CTCCTCATCATGCTAAGATGCAATGCCAAAATC |
| I228N | CTCCTCATCATGCTAAACTGCAATGCCAAAATC |
| I228D | CTCCTCATCATGCTAGACTGCAATGCCAAAATC |
| I228C | CTCCTCATCATGCTATGCTGCAATGCCAAAATC |
| I228Q | CTCCTCATCATGCTACAGTGCAATGCCAAAATC |
| I228E | CTCCTCATCATGCTAGAGTGCAATGCCAAAATC |
| I228G | CTCCTCATCATGCTAGGCTGCAATGCCAAAATC |
| I228H | CTCCTCATCATGCTACACTGCAATGCCAAAATC |
| I228L | CTCCTCATCATGCTACTGTGCAATGCCAAAATC |
| I228K | CTCCTCATCATGCTAAAGTGCAATGCCAAAATC |
| I228M | CTCCTCATCATGCTAATGTGCAATGCCAAAATC |
| I228F | CTCCTCATCATGCTATTCTGCAATGCCAAAATC |
| I228P | CTCCTCATCATGCTACCCTGCAATGCCAAAATC |
| I228S | CTCCTCATCATGCTAAGCTGCAATGCCAAAATC |
| I228T | CTCCTCATCATGCTAACCTGCAATGCCAAAATC |
| I228W | CTCCTCATCATGCTATGGTGCAATGCCAAAATC |
| I228Y | CTCCTCATCATGCTATACTGCAATGCCAAAATC |
| I228V | CTCCTCATCATGCTAGTGTGCAATGCCAAAATC |
| N230A | ATCATGCTAATCTGCGCCGCCAAAATCATCTTT |
| N230R | ATCATGCTAATCTGCAGAGCCAAAATCATCTTT |
| N230D | ATCATGCTAATCTGCGACGCCAAAATCATCTTT |

| | |
|---|---|
| N230C | ATCATGCTAATCTGCTGCGCCAAAATCATCTTT |
| N230Q | ATCATGCTAATCTGCCAGGCCAAAATCATCTTT |
| N230E | ATCATGCTAATCTGCGAGGCCAAAATCATCTTT |
| N230G | ATCATGCTAATCTGCGGCGCCAAAATCATCTTT |
| N230H | ATCATGCTAATCTGCCACGCCAAAATCATCTTT |
| N230I | ATCATGCTAATCTGCATCGCCAAAATCATCTTT |
| N230L | ATCATGCTAATCTGCCTGGCCAAAATCATCTTT |
| N230K | ATCATGCTAATCTGCAAGGCCAAAATCATCTTT |
| N230M | ATCATGCTAATCTGCATGGCCAAAATCATCTTT |
| N230F | ATCATGCTAATCTGCTTCGCCAAAATCATCTTT |
| N230P | ATCATGCTAATCTGCCCCGCCAAAATCATCTTT |
| N230S | ATCATGCTAATCTGCAGCGCCAAAATCATCTTT |
| N230T | ATCATGCTAATCTGCACCGCCAAAATCATCTTT |
| N230W | ATCATGCTAATCTGCTGGGCCAAAATCATCTTT |
| N230Y | ATCATGCTAATCTGCTACGCCAAAATCATCTTT |
| N230V | ATCATGCTAATCTGCGTGGCCAAAATCATCTTT |
| K232A | CTAATCTGCAATGCCGCCATCATCTTTGCTCTC |
| K232R | CTAATCTGCAATGCCAGAATCATCTTTGCTCTC |
| K232N | CTAATCTGCAATGCCAACATCATCTTTGCTCTC |
| K232D | CTAATCTGCAATGCCGACATCATCTTTGCTCTC |
| K232C | CTAATCTGCAATGCCTGCATCATCTTTGCTCTC |
| K232Q | CTAATCTGCAATGCCCAGATCATCTTTGCTCTC |
| K232E | CTAATCTGCAATGCCGAGATCATCTTTGCTCTC |
| K232G | CTAATCTGCAATGCCGGCATCATCTTTGCTCTC |
| K232H | CTAATCTGCAATGCCCACATCATCTTTGCTCTC |
| K232I | CTAATCTGCAATGCCATCATCATCTTTGCTCTC |
| K232L | CTAATCTGCAATGCCCTGATCATCTTTGCTCTC |
| K232M | CTAATCTGCAATGCCATGATCATCTTTGCTCTC |
| K232F | CTAATCTGCAATGCCTTCATCATCTTTGCTCTC |
| K232P | CTAATCTGCAATGCCCCCATCATCTTTGCTCTC |
| K232S | CTAATCTGCAATGCCAGCATCATCTTTGCTCTC |
| K232T | CTAATCTGCAATGCCACCATCATCTTTGCTCTC |
| K232W | CTAATCTGCAATGCCTGGATCATCTTTGCTCTC |
| K232Y | CTAATCTGCAATGCCTACATCATCTTTGCTCTC |
| K232V | CTAATCTGCAATGCCGTGATCATCTTTGCTCTC |
| T238A | ATCATCTTTGCTCTCGCCCGAGTCCTTCATCAA |
| T238R | ATCATCTTTGCTCTCAGACGAGTCCTTCATCAA |
| T238N | ATCATCTTTGCTCTCAACCGAGTCCTTCATCAA |
| T238D | ATCATCTTTGCTCTCGACCGAGTCCTTCATCAA |
| T238C | ATCATCTTTGCTCTCTGCCGAGTCCTTCATCAA |
| T238Q | ATCATCTTTGCTCTCCAGCGAGTCCTTCATCAA |

| | |
|---|---|
| T238E | ATCATCTTTGCTCTCGAGCGAGTCCTTCATCAA |
| T238G | ATCATCTTTGCTCTCGGCCGAGTCCTTCATCAA |
| T238H | ATCATCTTTGCTCTCCACCGAGTCCTTCATCAA |
| T238I | ATCATCTTTGCTCTCATCCGAGTCCTTCATCAA |
| T238L | ATCATCTTTGCTCTCCTGCGAGTCCTTCATCAA |
| T238K | ATCATCTTTGCTCTCAAGCGAGTCCTTCATCAA |
| T238M | ATCATCTTTGCTCTCATGCGAGTCCTTCATCAA |
| T238F | ATCATCTTTGCTCTCTTCCGAGTCCTTCATCAA |
| T238P | ATCATCTTTGCTCTCCCCCGAGTCCTTCATCAA |
| T238S | ATCATCTTTGCTCTCAGCCGAGTCCTTCATCAA |
| T238W | ATCATCTTTGCTCTCTGGCGAGTCCTTCATCAA |
| T238Y | ATCATCTTTGCTCTCTACCGAGTCCTTCATCAA |
| T238V | ATCATCTTTGCTCTCGTGCGAGTCCTTCATCAA |
| H242A | CTCACGCGAGTCCTTGCCCAAGACCCACGCAAA |
| H242R | CTCACGCGAGTCCTTAGACAAGACCCACGCAAA |
| H242N | CTCACGCGAGTCCTTAACCAAGACCCACGCAAA |
| H242D | CTCACGCGAGTCCTTGACCAAGACCCACGCAAA |
| H242C | CTCACGCGAGTCCTTTGCCAAGACCCACGCAAA |
| H242Q | CTCACGCGAGTCCTTCAGCAAGACCCACGCAAA |
| H242E | CTCACGCGAGTCCTTGAGCAAGACCCACGCAAA |
| H242G | CTCACGCGAGTCCTTGGCCAAGACCCACGCAAA |
| H242I | CTCACGCGAGTCCTTATCCAAGACCCACGCAAA |
| H242L | CTCACGCGAGTCCTTCTGCAAGACCCACGCAAA |
| H242K | CTCACGCGAGTCCTTAAGCAAGACCCACGCAAA |
| H242M | CTCACGCGAGTCCTTATGCAAGACCCACGCAAA |
| H242F | CTCACGCGAGTCCTTTTCCAAGACCCACGCAAA |
| H242P | CTCACGCGAGTCCTTCCCCAAGACCCACGCAAA |
| H242S | CTCACGCGAGTCCTTAGCCAAGACCCACGCAAA |
| H242T | CTCACGCGAGTCCTTACCCAAGACCCACGCAAA |
| H242W | CTCACGCGAGTCCTTTGGCAAGACCCACGCAAA |
| H242Y | CTCACGCGAGTCCTTTACCAAGACCCACGCAAA |
| H242V | CTCACGCGAGTCCTTGTGCAAGACCCACGCAAA |
| R246A | CTTCATCAAGACCCAGCCAAACTACAGCTGAAT |
| R246N | CTTCATCAAGACCCAAACAAACTACAGCTGAAT |
| R246D | CTTCATCAAGACCCAGACAAACTACAGCTGAAT |
| R246C | CTTCATCAAGACCCATGCAAACTACAGCTGAAT |
| R246Q | CTTCATCAAGACCCACAGAAACTACAGCTGAAT |
| R246E | CTTCATCAAGACCCAGAGAAACTACAGCTGAAT |
| R246G | CTTCATCAAGACCCAGGCAAACTACAGCTGAAT |
| R246H | CTTCATCAAGACCCACACAAACTACAGCTGAAT |
| R246I | CTTCATCAAGACCCAATCAAACTACAGCTGAAT |

| | |
|---|---|
| R246L | CTTCATCAAGACCCACTGAAACTACAGCTGAAT |
| R246K | CTTCATCAAGACCCAAAGAAACTACAGCTGAAT |
| R246M | CTTCATCAAGACCCAATGAAACTACAGCTGAAT |
| R246F | CTTCATCAAGACCCATTCAAACTACAGCTGAAT |
| R246P | CTTCATCAAGACCCACCCAAACTACAGCTGAAT |
| R246S | CTTCATCAAGACCCAAGCAAACTACAGCTGAAT |
| R246T | CTTCATCAAGACCCAACCAAACTACAGCTGAAT |
| R246W | CTTCATCAAGACCCATGGAAACTACAGCTGAAT |
| R246Y | CTTCATCAAGACCCATACAAACTACAGCTGAAT |
| R246V | CTTCATCAAGACCCAGTGAAACTACAGCTGAAT |
| L250A | CCACGCAAACTACAGGCCAATCAGTCCAAGAAT |
| L250R | CCACGCAAACTACAGAGAAATCAGTCCAAGAAT |
| L250N | CCACGCAAACTACAGAACAATCAGTCCAAGAAT |
| L250D | CCACGCAAACTACAGGACAATCAGTCCAAGAAT |
| L250C | CCACGCAAACTACAGTGCAATCAGTCCAAGAAT |
| L250Q | CCACGCAAACTACAGCAGAATCAGTCCAAGAAT |
| L250E | CCACGCAAACTACAGGAGAATCAGTCCAAGAAT |
| L250G | CCACGCAAACTACAGGGCAATCAGTCCAAGAAT |
| L250H | CCACGCAAACTACAGCACAATCAGTCCAAGAAT |
| L250I | CCACGCAAACTACAGATCAATCAGTCCAAGAAT |
| L250K | CCACGCAAACTACAGAAGAATCAGTCCAAGAAT |
| L250M | CCACGCAAACTACAGATGAATCAGTCCAAGAAT |
| L250F | CCACGCAAACTACAGTTCAATCAGTCCAAGAAT |
| L250P | CCACGCAAACTACAGCCCAATCAGTCCAAGAAT |
| L250S | CCACGCAAACTACAGAGCAATCAGTCCAAGAAT |
| L250T | CCACGCAAACTACAGACCAATCAGTCCAAGAAT |
| L250W | CCACGCAAACTACAGTGGAATCAGTCCAAGAAT |
| L250Y | CCACGCAAACTACAGTACAATCAGTCCAAGAAT |
| L250V | CCACGCAAACTACAGGTGAATCAGTCCAAGAAT |
| K254A | CAGCTGAATCAGTCCGCCAATAATATCCCAAGA |
| K254R | CAGCTGAATCAGTCCAGAAATAATATCCCAAGA |
| K254N | CAGCTGAATCAGTCCAACAATAATATCCCAAGA |
| K254D | CAGCTGAATCAGTCCGACAATAATATCCCAAGA |
| K254C | CAGCTGAATCAGTCCTGCAATAATATCCCAAGA |
| K254Q | CAGCTGAATCAGTCCCAGAATAATATCCCAAGA |
| K254E | CAGCTGAATCAGTCCGAGAATAATATCCCAAGA |
| K254G | CAGCTGAATCAGTCCGGCAATAATATCCCAAGA |
| K254H | CAGCTGAATCAGTCCCACAATAATATCCCAAGA |
| K254I | CAGCTGAATCAGTCCATCAATAATATCCCAAGA |
| K254L | CAGCTGAATCAGTCCCTGAATAATATCCCAAGA |
| K254M | CAGCTGAATCAGTCCATGAATAATATCCCAAGA |

| | |
|---|---|
| K254F | CAGCTGAATCAGTCCTTCAATAATATCCCAAGA |
| K254P | CAGCTGAATCAGTCCCCCAATAATATCCCAAGA |
| K254S | CAGCTGAATCAGTCCAGCAATAATATCCCAAGA |
| K254T | CAGCTGAATCAGTCCACCAATAATATCCCAAGA |
| K254W | CAGCTGAATCAGTCCTGGAATAATATCCCAAGA |
| K254Y | CAGCTGAATCAGTCCTACAATAATATCCCAAGA |
| K254V | CAGCTGAATCAGTCCGTGAATAATATCCCAAGA |
| P258A | TCCAAGAATAATATCGCCAGAGCTCGGCTGAGA |
| P258R | TCCAAGAATAATATCAGAAGAGCTCGGCTGAGA |
| P258N | TCCAAGAATAATATCAACAGAGCTCGGCTGAGA |
| P258D | TCCAAGAATAATATCGACAGAGCTCGGCTGAGA |
| P258C | TCCAAGAATAATATCTGCAGAGCTCGGCTGAGA |
| P258Q | TCCAAGAATAATATCCAGAGAGCTCGGCTGAGA |
| P258E | TCCAAGAATAATATCGAGAGAGCTCGGCTGAGA |
| P258G | TCCAAGAATAATATCGGCAGAGCTCGGCTGAGA |
| P258H | TCCAAGAATAATATCCACAGAGCTCGGCTGAGA |
| P258I | TCCAAGAATAATATCATCAGAGCTCGGCTGAGA |
| P258L | TCCAAGAATAATATCCTGAGAGCTCGGCTGAGA |
| P258K | TCCAAGAATAATATCAAGAGAGCTCGGCTGAGA |
| P258M | TCCAAGAATAATATCATGAGAGCTCGGCTGAGA |
| P258F | TCCAAGAATAATATCTTCAGAGCTCGGCTGAGA |
| P258S | TCCAAGAATAATATCAGCAGAGCTCGGCTGAGA |
| P258T | TCCAAGAATAATATCACCAGAGCTCGGCTGAGA |
| P258W | TCCAAGAATAATATCTGGAGAGCTCGGCTGAGA |
| P258Y | TCCAAGAATAATATCTACAGAGCTCGGCTGAGA |
| P258V | TCCAAGAATAATATCGTGAGAGCTCGGCTGAGA |
| L262A | ATCCCAAGAGCTCGGGCCAGAACGCTAAAGATG |
| L262R | ATCCCAAGAGCTCGGAGAAGAACGCTAAAGATG |
| L262N | ATCCCAAGAGCTCGGAACAGAACGCTAAAGATG |
| L262D | ATCCCAAGAGCTCGGGACAGAACGCTAAAGATG |
| L262C | ATCCCAAGAGCTCGGTGCAGAACGCTAAAGATG |
| L262Q | ATCCCAAGAGCTCGGCAGAGAACGCTAAAGATG |
| L262E | ATCCCAAGAGCTCGGGAGAGAACGCTAAAGATG |
| L262G | ATCCCAAGAGCTCGGGGCAGAACGCTAAAGATG |
| L262H | ATCCCAAGAGCTCGGCACAGAACGCTAAAGATG |
| L262I | ATCCCAAGAGCTCGGATCAGAACGCTAAAGATG |
| L262K | ATCCCAAGAGCTCGGAAGAGAACGCTAAAGATG |
| L262M | ATCCCAAGAGCTCGGATGAGAACGCTAAAGATG |
| L262F | ATCCCAAGAGCTCGGTTCAGAACGCTAAAGATG |
| L262P | ATCCCAAGAGCTCGGCCCAGAACGCTAAAGATG |
| L262S | ATCCCAAGAGCTCGGAGCAGAACGCTAAAGATG |

| | |
|---|---|
| L262T | ATCCCAAGAGCTCGGACCAGAACGCTAAAGATG |
| L262W | ATCCCAAGAGCTCGGTGGAGAACGCTAAAGATG |
| L262Y | ATCCCAAGAGCTCGGTACAGAACGCTAAAGATG |
| L262V | ATCCCAAGAGCTCGGGTGAGAACGCTAAAGATG |
| K266A | CGGCTGAGAACGCTAGCCATGACAGTCGCATTC |
| K266R | CGGCTGAGAACGCTAAGAATGACAGTCGCATTC |
| K266N | CGGCTGAGAACGCTAAACATGACAGTCGCATTC |
| K266D | CGGCTGAGAACGCTAGACATGACAGTCGCATTC |
| K266C | CGGCTGAGAACGCTATGCATGACAGTCGCATTC |
| K266Q | CGGCTGAGAACGCTACAGATGACAGTCGCATTC |
| K266E | CGGCTGAGAACGCTAGAGATGACAGTCGCATTC |
| K266G | CGGCTGAGAACGCTAGGCATGACAGTCGCATTC |
| K266H | CGGCTGAGAACGCTACACATGACAGTCGCATTC |
| K266I | CGGCTGAGAACGCTAATCATGACAGTCGCATTC |
| K266L | CGGCTGAGAACGCTACTGATGACAGTCGCATTC |
| K266M | CGGCTGAGAACGCTAATGATGACAGTCGCATTC |
| K266F | CGGCTGAGAACGCTATTCATGACAGTCGCATTC |
| K266P | CGGCTGAGAACGCTACCCATGACAGTCGCATTC |
| K266S | CGGCTGAGAACGCTAAGCATGACAGTCGCATTC |
| K266T | CGGCTGAGAACGCTAACCATGACAGTCGCATTC |
| K266W | CGGCTGAGAACGCTATGGATGACAGTCGCATTC |
| K266Y | CGGCTGAGAACGCTATACATGACAGTCGCATTC |
| K266V | CGGCTGAGAACGCTAGTGATGACAGTCGCATTC |
| F271A | AAGATGACAGTCGCAGCCGCTACCTCCTTTGTC |
| F271R | AAGATGACAGTCGCAAGAGCTACCTCCTTTGTC |
| F271N | AAGATGACAGTCGCAAACGCTACCTCCTTTGTC |
| F271D | AAGATGACAGTCGCAGACGCTACCTCCTTTGTC |
| F271C | AAGATGACAGTCGCATGCGCTACCTCCTTTGTC |
| F271Q | AAGATGACAGTCGCACAGGCTACCTCCTTTGTC |
| F271E | AAGATGACAGTCGCAGAGGCTACCTCCTTTGTC |
| F271G | AAGATGACAGTCGCAGGCGCTACCTCCTTTGTC |
| F271H | AAGATGACAGTCGCACACGCTACCTCCTTTGTC |
| F271I | AAGATGACAGTCGCAATCGCTACCTCCTTTGTC |
| F271L | AAGATGACAGTCGCACTGGCTACCTCCTTTGTC |
| F271K | AAGATGACAGTCGCAAAGGCTACCTCCTTTGTC |
| F271M | AAGATGACAGTCGCAATGGCTACCTCCTTTGTC |
| F271P | AAGATGACAGTCGCACCCGCTACCTCCTTTGTC |
| F271S | AAGATGACAGTCGCAAGCGCTACCTCCTTTGTC |
| F271T | AAGATGACAGTCGCAACCGCTACCTCCTTTGTC |
| F271W | AAGATGACAGTCGCATGGGCTACCTCCTTTGTC |
| F271Y | AAGATGACAGTCGCATACGCTACCTCCTTTGTC |

| | |
|---|---|
| F271V | AAGATGACAGTCGCAGTGGCTACCTCCTTTGTC |
| T273A | ACAGTCGCATTCGCTGCCTCCTTTGTCGTCTGC |
| T273R | ACAGTCGCATTCGCTAGATCCTTTGTCGTCTGC |
| T273N | ACAGTCGCATTCGCTAACTCCTTTGTCGTCTGC |
| T273D | ACAGTCGCATTCGCTGACTCCTTTGTCGTCTGC |
| T273C | ACAGTCGCATTCGCTTGCTCCTTTGTCGTCTGC |
| T273Q | ACAGTCGCATTCGCTCAGTCCTTTGTCGTCTGC |
| T273E | ACAGTCGCATTCGCTGAGTCCTTTGTCGTCTGC |
| T273G | ACAGTCGCATTCGCTGGCTCCTTTGTCGTCTGC |
| T273H | ACAGTCGCATTCGCTCACTCCTTTGTCGTCTGC |
| T273I | ACAGTCGCATTCGCTATCTCCTTTGTCGTCTGC |
| T273L | ACAGTCGCATTCGCTCTGTCCTTTGTCGTCTGC |
| T273K | ACAGTCGCATTCGCTAAGTCCTTTGTCGTCTGC |
| T273M | ACAGTCGCATTCGCTATGTCCTTTGTCGTCTGC |
| T273F | ACAGTCGCATTCGCTTTCTCCTTTGTCGTCTGC |
| T273P | ACAGTCGCATTCGCTCCCTCCTTTGTCGTCTGC |
| T273S | ACAGTCGCATTCGCTAGCTCCTTTGTCGTCTGC |
| T273W | ACAGTCGCATTCGCTTGGTCCTTTGTCGTCTGC |
| T273Y | ACAGTCGCATTCGCTTACTCCTTTGTCGTCTGC |
| T273V | ACAGTCGCATTCGCTGTGTCCTTTGTCGTCTGC |
| F275A | GCATTCGCTACCTCCGCCGTCGTCTGCTGGACT |
| F275R | GCATTCGCTACCTCCAGAGTCGTCTGCTGGACT |
| F275N | GCATTCGCTACCTCCAACGTCGTCTGCTGGACT |
| F275D | GCATTCGCTACCTCCGACGTCGTCTGCTGGACT |
| F275C | GCATTCGCTACCTCCTGCGTCGTCTGCTGGACT |
| F275Q | GCATTCGCTACCTCCCAGGTCGTCTGCTGGACT |
| F275E | GCATTCGCTACCTCCGAGGTCGTCTGCTGGACT |
| F275G | GCATTCGCTACCTCCGGCGTCGTCTGCTGGACT |
| F275H | GCATTCGCTACCTCCCACGTCGTCTGCTGGACT |
| F275I | GCATTCGCTACCTCCATCGTCGTCTGCTGGACT |
| F275L | GCATTCGCTACCTCCCTGGTCGTCTGCTGGACT |
| F275K | GCATTCGCTACCTCCAAGGTCGTCTGCTGGACT |
| F275M | GCATTCGCTACCTCCATGGTCGTCTGCTGGACT |
| F275P | GCATTCGCTACCTCCCCCGTCGTCTGCTGGACT |
| F275S | GCATTCGCTACCTCCAGCGTCGTCTGCTGGACT |
| F275T | GCATTCGCTACCTCCACCGTCGTCTGCTGGACT |
| F275W | GCATTCGCTACCTCCTGGGTCGTCTGCTGGACT |
| F275Y | GCATTCGCTACCTCCTACGTCGTCTGCTGGACT |
| F275V | GCATTCGCTACCTCCGTGGTCGTCTGCTGGACT |
| V277A | GCTACCTCCTTTGTCGCCTGCTGGACTCCCTAC |
| V277R | GCTACCTCCTTTGTCAGATGCTGGACTCCCTAC |

| | |
|---|---|
| V277N | GCTACCTCCTTTGTCAACTGCTGGACTCCCTAC |
| V277D | GCTACCTCCTTTGTCGACTGCTGGACTCCCTAC |
| V277C | GCTACCTCCTTTGTCTGCTGCTGGACTCCCTAC |
| V277Q | GCTACCTCCTTTGTCCAGTGCTGGACTCCCTAC |
| V277E | GCTACCTCCTTTGTCGAGTGCTGGACTCCCTAC |
| V277G | GCTACCTCCTTTGTCGGCTGCTGGACTCCCTAC |
| V277H | GCTACCTCCTTTGTCCACTGCTGGACTCCCTAC |
| V277I | GCTACCTCCTTTGTCATCTGCTGGACTCCCTAC |
| V277L | GCTACCTCCTTTGTCCTGTGCTGGACTCCCTAC |
| V277K | GCTACCTCCTTTGTCAAGTGCTGGACTCCCTAC |
| V277M | GCTACCTCCTTTGTCATGTGCTGGACTCCCTAC |
| V277F | GCTACCTCCTTTGTCTTCTGCTGGACTCCCTAC |
| V277P | GCTACCTCCTTTGTCCCCTGCTGGACTCCCTAC |
| V277S | GCTACCTCCTTTGTCAGCTGCTGGACTCCCTAC |
| V277T | GCTACCTCCTTTGTCACCTGCTGGACTCCCTAC |
| V277W | GCTACCTCCTTTGTCTGGTGCTGGACTCCCTAC |
| V277Y | GCTACCTCCTTTGTCTACTGCTGGACTCCCTAC |
| C278A | ACCTCCTTTGTCGTCGCCTGGACTCCCTACTAT |
| C278R | ACCTCCTTTGTCGTCAGATGGACTCCCTACTAT |
| C278N | ACCTCCTTTGTCGTCAACTGGACTCCCTACTAT |
| C278D | ACCTCCTTTGTCGTCGACTGGACTCCCTACTAT |
| C278Q | ACCTCCTTTGTCGTCCAGTGGACTCCCTACTAT |
| C278E | ACCTCCTTTGTCGTCGAGTGGACTCCCTACTAT |
| C278G | ACCTCCTTTGTCGTCGGCTGGACTCCCTACTAT |
| C278H | ACCTCCTTTGTCGTCCACTGGACTCCCTACTAT |
| C278I | ACCTCCTTTGTCGTCATCTGGACTCCCTACTAT |
| C278L | ACCTCCTTTGTCGTCCTGTGGACTCCCTACTAT |
| C278K | ACCTCCTTTGTCGTCAAGTGGACTCCCTACTAT |
| C278M | ACCTCCTTTGTCGTCATGTGGACTCCCTACTAT |
| C278F | ACCTCCTTTGTCGTCTTCTGGACTCCCTACTAT |
| C278P | ACCTCCTTTGTCGTCCCCTGGACTCCCTACTAT |
| C278S | ACCTCCTTTGTCGTCAGCTGGACTCCCTACTAT |
| C278T | ACCTCCTTTGTCGTCACCTGGACTCCCTACTAT |
| C278W | ACCTCCTTTGTCGTCTGGTGGACTCCCTACTAT |
| C278Y | ACCTCCTTTGTCGTCTACTGGACTCCCTACTAT |
| C278V | ACCTCCTTTGTCGTCGTGTGGACTCCCTACTAT |
| L285A | ACTCCCTACTATGTCGCCGGCATTTGGTACTGG |
| L285R | ACTCCCTACTATGTCAGAGGCATTTGGTACTGG |
| L285N | ACTCCCTACTATGTCAACGGCATTTGGTACTGG |
| L285D | ACTCCCTACTATGTCGACGGCATTTGGTACTGG |
| L285C | ACTCCCTACTATGTCTGCGGCATTTGGTACTGG |

| | |
|---|---|
| L285Q | ACTCCCTACTATGTCCAGGGCATTTGGTACTGG |
| L285E | ACTCCCTACTATGTCGAGGGCATTTGGTACTGG |
| L285G | ACTCCCTACTATGTCGGCGGCATTTGGTACTGG |
| L285H | ACTCCCTACTATGTCCACGGCATTTGGTACTGG |
| L285I | ACTCCCTACTATGTCATCGGCATTTGGTACTGG |
| L285K | ACTCCCTACTATGTCAAGGGCATTTGGTACTGG |
| L285M | ACTCCCTACTATGTCATGGGCATTTGGTACTGG |
| L285F | ACTCCCTACTATGTCTTCGGCATTTGGTACTGG |
| L285P | ACTCCCTACTATGTCCCCGGCATTTGGTACTGG |
| L285S | ACTCCCTACTATGTCAGCGGCATTTGGTACTGG |
| L285T | ACTCCCTACTATGTCACCGGCATTTGGTACTGG |
| L285W | ACTCCCTACTATGTCTGGGGCATTTGGTACTGG |
| L285Y | ACTCCCTACTATGTCTACGGCATTTGGTACTGG |
| L285V | ACTCCCTACTATGTCGTGGGCATTTGGTACTGG |
| W288A | TATGTCCTAGGCATTGCCTACTGGTTTGATCCA |
| W288R | TATGTCCTAGGCATTAGATACTGGTTTGATCCA |
| W288N | TATGTCCTAGGCATTAACTACTGGTTTGATCCA |
| W288D | TATGTCCTAGGCATTGACTACTGGTTTGATCCA |
| W288C | TATGTCCTAGGCATTTGCTACTGGTTTGATCCA |
| W288Q | TATGTCCTAGGCATTCAGTACTGGTTTGATCCA |
| W288E | TATGTCCTAGGCATTGAGTACTGGTTTGATCCA |
| W288G | TATGTCCTAGGCATTGGCTACTGGTTTGATCCA |
| W288H | TATGTCCTAGGCATTCACTACTGGTTTGATCCA |
| W288I | TATGTCCTAGGCATTATCTACTGGTTTGATCCA |
| W288L | TATGTCCTAGGCATTCTGTACTGGTTTGATCCA |
| W288K | TATGTCCTAGGCATTAAGTACTGGTTTGATCCA |
| W288M | TATGTCCTAGGCATTATGTACTGGTTTGATCCA |
| W288F | TATGTCCTAGGCATTTTCTACTGGTTTGATCCA |
| W288P | TATGTCCTAGGCATTCCCTACTGGTTTGATCCA |
| W288S | TATGTCCTAGGCATTAGCTACTGGTTTGATCCA |
| W288T | TATGTCCTAGGCATTACCTACTGGTTTGATCCA |
| W288Y | TATGTCCTAGGCATTTACTACTGGTTTGATCCA |
| W288V | TATGTCCTAGGCATTGTGTACTGGTTTGATCCA |
| M295A | TGGTTTGATCCAGAAGCCTTGAACAGGGTGTCA |
| M295R | TGGTTTGATCCAGAAAGATTGAACAGGGTGTCA |
| M295N | TGGTTTGATCCAGAAAACTTGAACAGGGTGTCA |
| M295D | TGGTTTGATCCAGAAGACTTGAACAGGGTGTCA |
| M295C | TGGTTTGATCCAGAATGCTTGAACAGGGTGTCA |
| M295Q | TGGTTTGATCCAGAACAGTTGAACAGGGTGTCA |
| M295E | TGGTTTGATCCAGAAGAGTTGAACAGGGTGTCA |
| M295G | TGGTTTGATCCAGAAGGCTTGAACAGGGTGTCA |

| | |
|---|---|
| M295H | TGGTTTGATCCAGAACACTTGAACAGGGTGTCA |
| M295I | TGGTTTGATCCAGAAATCTTGAACAGGGTGTCA |
| M295L | TGGTTTGATCCAGAACTGTTGAACAGGGTGTCA |
| M295K | TGGTTTGATCCAGAAAAGTTGAACAGGGTGTCA |
| M295F | TGGTTTGATCCAGAATTCTTGAACAGGGTGTCA |
| M295P | TGGTTTGATCCAGAACCCTTGAACAGGGTGTCA |
| M295S | TGGTTTGATCCAGAAAGCTTGAACAGGGTGTCA |
| M295T | TGGTTTGATCCAGAAACCTTGAACAGGGTGTCA |
| M295W | TGGTTTGATCCAGAATGGTTGAACAGGGTGTCA |
| M295Y | TGGTTTGATCCAGAATACTTGAACAGGGTGTCA |
| M295V | TGGTTTGATCCAGAAGTGTTGAACAGGGTGTCA |
| V299A | GAAATGTTGAACAGGGCCTCAGAGCCAGTGAAT |
| V299R | GAAATGTTGAACAGGAGATCAGAGCCAGTGAAT |
| V299N | GAAATGTTGAACAGGAACTCAGAGCCAGTGAAT |
| V299D | GAAATGTTGAACAGGGACTCAGAGCCAGTGAAT |
| V299C | GAAATGTTGAACAGGTGCTCAGAGCCAGTGAAT |
| V299Q | GAAATGTTGAACAGGCAGTCAGAGCCAGTGAAT |
| V299E | GAAATGTTGAACAGGGAGTCAGAGCCAGTGAAT |
| V299G | GAAATGTTGAACAGGGGCTCAGAGCCAGTGAAT |
| V299H | GAAATGTTGAACAGGCACTCAGAGCCAGTGAAT |
| V299I | GAAATGTTGAACAGGATCTCAGAGCCAGTGAAT |
| V299L | GAAATGTTGAACAGGCTGTCAGAGCCAGTGAAT |
| V299K | GAAATGTTGAACAGGAAGTCAGAGCCAGTGAAT |
| V299M | GAAATGTTGAACAGGATGTCAGAGCCAGTGAAT |
| V299F | GAAATGTTGAACAGGTTCTCAGAGCCAGTGAAT |
| V299P | GAAATGTTGAACAGGCCCTCAGAGCCAGTGAAT |
| V299S | GAAATGTTGAACAGGAGCTCAGAGCCAGTGAAT |
| V299T | GAAATGTTGAACAGGACCTCAGAGCCAGTGAAT |
| V299W | GAAATGTTGAACAGGTGGTCAGAGCCAGTGAAT |
| V299Y | GAAATGTTGAACAGGTACTCAGAGCCAGTGAAT |
| V303A | AGGGTGTCAGAGCCAGCCAATCACTTTTTCTTT |
| V303R | AGGGTGTCAGAGCCAAGAAATCACTTTTTCTTT |
| V303N | AGGGTGTCAGAGCCAAACAATCACTTTTTCTTT |
| V303D | AGGGTGTCAGAGCCAGACAATCACTTTTTCTTT |
| V303C | AGGGTGTCAGAGCCATGCAATCACTTTTTCTTT |
| V303Q | AGGGTGTCAGAGCCACAGAATCACTTTTTCTTT |
| V303E | AGGGTGTCAGAGCCAGAGAATCACTTTTTCTTT |
| V303G | AGGGTGTCAGAGCCAGGCAATCACTTTTTCTTT |
| V303H | AGGGTGTCAGAGCCACACAATCACTTTTTCTTT |
| V303I | AGGGTGTCAGAGCCAATCAATCACTTTTTCTTT |
| V303L | AGGGTGTCAGAGCCACTGAATCACTTTTTCTTT |

| | |
|---|---|
| V303K | AGGGTGTCAGAGCCAAAGAATCACTTTTTCTTT |
| V303M | AGGGTGTCAGAGCCAATGAATCACTTTTTCTTT |
| V303F | AGGGTGTCAGAGCCATTCAATCACTTTTTCTTT |
| V303P | AGGGTGTCAGAGCCACCCAATCACTTTTTCTTT |
| V303S | AGGGTGTCAGAGCCAAGCAATCACTTTTTCTTT |
| V303T | AGGGTGTCAGAGCCAACCAATCACTTTTTCTTT |
| V303W | AGGGTGTCAGAGCCATGGAATCACTTTTTCTTT |
| V303Y | AGGGTGTCAGAGCCATACAATCACTTTTTCTTT |
| F310A | CACTTTTTCTTTCTCGCCGCTTTCCTAAACCCG |
| F310R | CACTTTTTCTTTCTCAGAGCTTTCCTAAACCCG |
| F310N | CACTTTTTCTTTCTCAACGCTTTCCTAAACCCG |
| F310D | CACTTTTTCTTTCTCGACGCTTTCCTAAACCCG |
| F310C | CACTTTTTCTTTCTCTGCGCTTTCCTAAACCCG |
| F310Q | CACTTTTTCTTTCTCCAGGCTTTCCTAAACCCG |
| F310E | CACTTTTTCTTTCTCGAGGCTTTCCTAAACCCG |
| F310G | CACTTTTTCTTTCTCGGCGCTTTCCTAAACCCG |
| F310H | CACTTTTTCTTTCTCCACGCTTTCCTAAACCCG |
| F310I | CACTTTTTCTTTCTCATCGCTTTCCTAAACCCG |
| F310L | CACTTTTTCTTTCTCCTGGCTTTCCTAAACCCG |
| F310K | CACTTTTTCTTTCTCAAGGCTTTCCTAAACCCG |
| F310M | CACTTTTTCTTTCTCATGGCTTTCCTAAACCCG |
| F310P | CACTTTTTCTTTCTCCCCGCTTTCCTAAACCCG |
| F310S | CACTTTTTCTTTCTCAGCGCTTTCCTAAACCCG |
| F310T | CACTTTTTCTTTCTCACCGCTTTCCTAAACCCG |
| F310W | CACTTTTTCTTTCTCTGGGCTTTCCTAAACCCG |
| F310Y | CACTTTTTCTTTCTCTACGCTTTCCTAAACCCG |
| F310V | CACTTTTTCTTTCTCGTGGCTTTCCTAAACCCG |
| A311R | TTTTTCTTTCTCTTTAGATTCCTAAACCCGTGC |
| A311N | TTTTTCTTTCTCTTTAACTTCCTAAACCCGTGC |
| A311D | TTTTTCTTTCTCTTTGACTTCCTAAACCCGTGC |
| A311C | TTTTTCTTTCTCTTTTGCTTCCTAAACCCGTGC |
| A311Q | TTTTTCTTTCTCTTTCAGTTCCTAAACCCGTGC |
| A311E | TTTTTCTTTCTCTTTGAGTTCCTAAACCCGTGC |
| A311G | TTTTTCTTTCTCTTTGGCTTCCTAAACCCGTGC |
| A311H | TTTTTCTTTCTCTTTCACTTCCTAAACCCGTGC |
| A311I | TTTTTCTTTCTCTTTATCTTCCTAAACCCGTGC |
| A311L | TTTTTCTTTCTCTTTCTGTTCCTAAACCCGTGC |
| A311K | TTTTTCTTTCTCTTTAAGTTCCTAAACCCGTGC |
| A311M | TTTTTCTTTCTCTTTATGTTCCTAAACCCGTGC |
| A311F | TTTTTCTTTCTCTTTTTCTTCCTAAACCCGTGC |
| A311P | TTTTTCTTTCTCTTTCCCTTCCTAAACCCGTGC |

| | |
|---|---|
| A311S | TTTTTCTTTCTCTTTAGCTTCCTAAACCCGTGC |
| A311T | TTTTTCTTTCTCTTTACCTTCCTAAACCCGTGC |
| A311W | TTTTTCTTTCTCTTTTGGTTCCTAAACCCGTGC |
| A311Y | TTTTTCTTTCTCTTTTACTTCCTAAACCCGTGC |
| A311V | TTTTTCTTTCTCTTTGTGTTCCTAAACCCGTGC |
| F317A | TTCCTAAACCCGTGCGCCGACCCACTCATATAT |
| F317R | TTCCTAAACCCGTGCAGAGACCCACTCATATAT |
| F317N | TTCCTAAACCCGTGCAACGACCCACTCATATAT |
| F317D | TTCCTAAACCCGTGCGACGACCCACTCATATAT |
| F317C | TTCCTAAACCCGTGCTGCGACCCACTCATATAT |
| F317Q | TTCCTAAACCCGTGCCAGGACCCACTCATATAT |
| F317E | TTCCTAAACCCGTGCGAGGACCCACTCATATAT |
| F317G | TTCCTAAACCCGTGCGGCGACCCACTCATATAT |
| F317H | TTCCTAAACCCGTGCCACGACCCACTCATATAT |
| F317I | TTCCTAAACCCGTGCATCGACCCACTCATATAT |
| F317L | TTCCTAAACCCGTGCCTGGACCCACTCATATAT |
| F317K | TTCCTAAACCCGTGCAAGGACCCACTCATATAT |
| F317M | TTCCTAAACCCGTGCATGGACCCACTCATATAT |
| F317P | TTCCTAAACCCGTGCCCCGACCCACTCATATAT |
| F317S | TTCCTAAACCCGTGCAGCGACCCACTCATATAT |
| F317T | TTCCTAAACCCGTGCACCGACCCACTCATATAT |
| F317W | TTCCTAAACCCGTGCTGGGACCCACTCATATAT |
| F317Y | TTCCTAAACCCGTGCTACGACCCACTCATATAT |
| F317V | TTCCTAAACCCGTGCGTGGACCCACTCATATAT |
| D318A | CTAAACCCGTGCTTCGCCCCACTCATATATGGG |
| D318R | CTAAACCCGTGCTTCAGACCACTCATATATGGG |
| D318N | CTAAACCCGTGCTTCAACCCACTCATATATGGG |
| D318C | CTAAACCCGTGCTTCTGCCCACTCATATATGGG |
| D318Q | CTAAACCCGTGCTTCCAGCCACTCATATATGGG |
| D318E | CTAAACCCGTGCTTCGAGCCACTCATATATGGG |
| D318G | CTAAACCCGTGCTTCGGCCCACTCATATATGGG |
| D318H | CTAAACCCGTGCTTCCACCCACTCATATATGGG |
| D318I | CTAAACCCGTGCTTCATCCCACTCATATATGGG |
| D318L | CTAAACCCGTGCTTCCTGCCACTCATATATGGG |
| D318K | CTAAACCCGTGCTTCAAGCCACTCATATATGGG |
| D318M | CTAAACCCGTGCTTCATGCCACTCATATATGGG |
| D318F | CTAAACCCGTGCTTCTTCCCACTCATATATGGG |
| D318P | CTAAACCCGTGCTTCCCCCCACTCATATATGGG |
| D318S | CTAAACCCGTGCTTCAGCCCACTCATATATGGG |
| D318T | CTAAACCCGTGCTTCACCCCACTCATATATGGG |
| D318W | CTAAACCCGTGCTTCTGGCCACTCATATATGGG |

| | |
|---|---|
| D318Y | CTAAACCCGTGCTTCTACCCACTCATATATGGG |
| D318V | CTAAACCCGTGCTTCGTGCCACTCATATATGGG |
| L320A | CCGTGCTTCGACCCAGCCATATATGGGTATTTC |
| L320R | CCGTGCTTCGACCCAAGAATATATGGGTATTTC |
| L320N | CCGTGCTTCGACCCAAACATATATGGGTATTTC |
| L320D | CCGTGCTTCGACCCAGACATATATGGGTATTTC |
| L320C | CCGTGCTTCGACCCATGCATATATGGGTATTTC |
| L320Q | CCGTGCTTCGACCCACAGATATATGGGTATTTC |
| L320E | CCGTGCTTCGACCCAGAGATATATGGGTATTTC |
| L320G | CCGTGCTTCGACCCAGGCATATATGGGTATTTC |
| L320H | CCGTGCTTCGACCCACACATATATGGGTATTTC |
| L320I | CCGTGCTTCGACCCAATCATATATGGGTATTTC |
| L320K | CCGTGCTTCGACCCAAAGATATATGGGTATTTC |
| L320M | CCGTGCTTCGACCCAATGATATATGGGTATTTC |
| L320F | CCGTGCTTCGACCCATTCATATATGGGTATTTC |
| L320P | CCGTGCTTCGACCCACCCATATATGGGTATTTC |
| L320S | CCGTGCTTCGACCCAAGCATATATGGGTATTTC |
| L320T | CCGTGCTTCGACCCAACCATATATGGGTATTTC |
| L320W | CCGTGCTTCGACCCATGGATATATGGGTATTTC |
| L320Y | CCGTGCTTCGACCCATACATATATGGGTATTTC |
| L320V | CCGTGCTTCGACCCAGTGATATATGGGTATTTC |
| Y322A | TTCGACCCACTCATAGCCGGGTATTTCTCTTTG |
| Y322R | TTCGACCCACTCATAAGAGGGTATTTCTCTTTG |
| Y322N | TTCGACCCACTCATAAACGGGTATTTCTCTTTG |
| Y322D | TTCGACCCACTCATAGACGGGTATTTCTCTTTG |
| Y322C | TTCGACCCACTCATATGCGGGTATTTCTCTTTG |
| Y322Q | TTCGACCCACTCATACAGGGGTATTTCTCTTTG |
| Y322E | TTCGACCCACTCATAGAGGGGTATTTCTCTTTG |
| Y322G | TTCGACCCACTCATAGGCGGGTATTTCTCTTTG |
| Y322H | TTCGACCCACTCATACACGGGTATTTCTCTTTG |
| Y322I | TTCGACCCACTCATAATCGGGTATTTCTCTTTG |
| Y322L | TTCGACCCACTCATACTGGGGTATTTCTCTTTG |
| Y322K | TTCGACCCACTCATAAAGGGGTATTTCTCTTTG |
| Y322M | TTCGACCCACTCATAATGGGGTATTTCTCTTTG |
| Y322F | TTCGACCCACTCATATTCGGGTATTTCTCTTTG |
| Y322P | TTCGACCCACTCATACCCGGGTATTTCTCTTTG |
| Y322S | TTCGACCCACTCATAAGCGGGTATTTCTCTTTG |
| Y322T | TTCGACCCACTCATAACCGGGTATTTCTCTTTG |
| Y322W | TTCGACCCACTCATATGGGGGTATTTCTCTTTG |
| Y322V | TTCGACCCACTCATAGTGGGGTATTTCTCTTTG |
| L327A | TATGGGTATTTCTCTGCCTAGGGCGGCCGCTCG |

| | |
|---|---|
| L327R | TATGGGTATTTCTCTAGATAGGGCGGCCGCTCG |
| L327N | TATGGGTATTTCTCTAACTAGGGCGGCCGCTCG |
| L327D | TATGGGTATTTCTCTGACTAGGGCGGCCGCTCG |
| L327C | TATGGGTATTTCTCTTGCTAGGGCGGCCGCTCG |
| L327Q | TATGGGTATTTCTCTCAGTAGGGCGGCCGCTCG |
| L327E | TATGGGTATTTCTCTGAGTAGGGCGGCCGCTCG |
| L327G | TATGGGTATTTCTCTGGCTAGGGCGGCCGCTCG |
| L327H | TATGGGTATTTCTCTCACTAGGGCGGCCGCTCG |
| L327I | TATGGGTATTTCTCTATCTAGGGCGGCCGCTCG |
| L327K | TATGGGTATTTCTCTAAGTAGGGCGGCCGCTCG |
| L327M | TATGGGTATTTCTCTATGTAGGGCGGCCGCTCG |
| L327F | TATGGGTATTTCTCTTTCTAGGGCGGCCGCTCG |
| L327P | TATGGGTATTTCTCTCCCTAGGGCGGCCGCTCG |
| L327S | TATGGGTATTTCTCTAGCTAGGGCGGCCGCTCG |
| L327T | TATGGGTATTTCTCTACCTAGGGCGGCCGCTCG |
| L327W | TATGGGTATTTCTCTTGGTAGGGCGGCCGCTCG |
| L327Y | TATGGGTATTTCTCTTACTAGGGCGGCCGCTCG |
| L327V | TATGGGTATTTCTCTGTGTAGGGCGGCCGCTCG |
